# Multivariate distributions of behavioural, morphological, and ontogenetic traits in hybrids bring new insights into the divergence of sympatric Arctic charr morphs

**DOI:** 10.1101/2020.10.14.339911

**Authors:** Quentin J.B. Horta-Lacueva, Sigurður S. Snorrason, Michael B. Morrissey, Camille A. Leblanc, Kalina H. Kapralova

## Abstract

Studying the development of fitness related traits in hybrids from populations diverging in sympatry is a fundamental approach to understand the processes of speciation. However, such traits are often affected by covariance structures that complicate the comprehension of these processes, especially because the interactive relationships between traits of different nature (e.g. morphology, behaviour, life-history) remain largely unknown in this context. In a common garden setup, we conducted an extensive examination of phenotypic traits suspected to be involved in the divergence of two recently evolved morphs of Arctic charr (*Salvelinus alpinus*), and investigated the consequences of potential patterns of trait covariance on the phenotype of their hybrids. We observed differences among morphs in overall phenotypic variance and in trait correlations. Phenotypic contrainsts also tended to be reduced in the hybrids, which corroborates the narrative of hybridization facilitating adaptive divergence by relaxing trait covariance. However, the hybrids were associated with reduced phenotypic variance at different scales (i.e. at the scale of the entire **P** matrix and in different parts of the multivariate space), and we identified stronger correlations between several ontogenetic and morphological traits in the hybrids than in both morphs. These findings suggest a limited potential for hybridization to generate phenotypic novelty, and emphasise the need for multivariate approaches conciliating ontogenetic, morphological and behavioural processes to study the processes of adaptive divergence and speciation.

## Introduction

Understanding how phenotypic traits subjected to divergent selection evolve is essential to comprehend the processes of adaptive divergence and speciation [2–5]. Fitness-based approaches are valuable ways to study how reproductive isolation emerges as a by-product of divergent selection between populations facing contrasting ecological conditions (e.g. following the colonisation of new niches) [3,6,7]. In this context, reproductive isolation often relates to reduced fitness in hybrids whose values for specific traits under divergent selection are intermediate or fall outside of the range of parental values (i.e. transgressive characters) [8–10]. However, traits are rarely independent entities and the responses to selection are influenced by their interactions with one-another because of functional trade-offs [11,12] and genetic constraints like pleiotropy and linkage disequilibrium [13,14]. Adaptive evolution may also be complicated by the effects of correlational selection, through which fitness is determined by optimal combinations of multiple traits [15,16]. While these evolutionary aspects have long been studied in the field of quantitative genetics, and while classical models of ecological speciation are based on the effects of pleiotropy and/or of large sets of co-selected genes [3,17,18], our understanding of how trait covariance may affect the phenotypic differentiation of populations and the associated build-up of reproductive isolation remains limited.

The processes of adaptive divergence and speciation are, however, often studied by focusing on one or a limited number of traits of the same nature, most often related to morphology and to some extent to physiology and behaviour [3] (but see [19] for an integrative study on life-history and morphology). Further difficulties in understanding these processes emerge as the development of characters involved in population divergence can be subjected to interactions among a diversity of traits encompassing life-history, development [20], and (in animals) behavioural processes [21,22]. In teleost fish and amphibians, for example, early developmental and broad-sense life-history traits (e.g. yolk-sac resorption, growth rate, onset of first feeding) interact with one another through mechanisms like energy allocation trade-offs [23–26], and some of these traits influence the establishment of behaviours appearing later in life (e.g. foraging activity, anti-predator response) [23–28]. Moreover, variations in feeding behaviour can affect the development of trophic morphology [29,30] and is intertwined with life-history traits through energy acquisition and allocation [31–33].

The implications of trait covariances on the processes of adaptive divergence and speciation remain relatively unexplored. Such covariances may on the one hand constrain divergence but could on the other hand be powerful fine-tuners of the development of adaptive phenotypes (Figure S1). Very little is known about how these mechanisms relate to the emergence of reproductive isolation, especially regarding the development of hybrid phenotypes [3]. Simulations have shown that genetic constraints like pleiotropy can limit the development of transgressive hybrid traits to varying extents, depending on different bioenergetic and evolutionary factors [34]. However, empirical evidences indicate that hybridization can break down such genetic constraints, thus facilitating the development of transgressive hybrid traits [9,35]. It is therefore critical to understand how trait correlations influence the development of diverging phenotypes and those of their hybrids before embarking into studying their proximate mechanisms (e.g. genomic architecture, resource allocation trade-offs) and their evolutionary consequences (e.g. selection against hybrids as a reproductive barrier).

Polymorphic fish from Northern freshwater lakes are particularly well-suited models to study the processes of phenotypic divergence [36]. The evolution of these fish fits the narrative of resource polymorphism, through which different forms (*i*.*e*. morphs) have emerged from ancestral populations that invaded multiple, unoccupied niches within the same geographical system [31]. Such diversification often follows the colonisation of deglaciated lakes, where the diverging morphs (generally segregating between benthic and pelagic habitats) differ in morphology, life-history traits and/or behaviour [37,38]. Various levels of reproductive isolation are encountered among these systems, ranging from single populations with continuous variation, to discrete varieties with more-or-less reversible reproductive barriers, to completely reproductively isolated species [8,39,40]. In recent years a growing number of cases have been reported where post-glacial morphs are found (at least in their current state) in sympatry [40–43]. These geographical and evolutionary systems facilitate the explorations of the mechanisms of adaptive divergence and speciation because of the reduced confounding effects of long and complex evolutionary histories [44].

Using multivariate phenotypic data on morphology, behaviour and ontogeny, and considering different developmental stages, we characterized phenotypic variations among two of the four sympatric morphs of Arctic charr (*Salvelinus alpinus*) from lake Thingvallavatn, Iceland, and of their hybrids. These morphs are the small-benthic (SB) and the planktivorous charr (PL), which constitute two genetically differentiated populations [45–47] and differ in head and body shape, habitat use, diet, life-history and parasites [48–50]. The SB charr live in the interstitial spaces of a lava matrix forming the stony littoral zone of the lake, where they forage on benthic invertebrates. The PL charr utilizes the pelagic zone of the lake where they feed on zooplankton and emerging chironomids. Because these two habitats differ extensively in their physical and ecological characteristics [50,51], the different selective regimes experienced by each morph are expected to affect a wide variety to traits. Previous studies already indicate that the PL and the SB charr have evolved genetically based differences in their embryonic growth [49], craniofacial development [52,53], and foraging strategy [54]. The two morphs overlap in their spawning time and places [55] but recent estimates of gene flow indicate substantial reproductive isolation [46,47]. Fertile hybrids (at least of the generation F_1_) can however easily be produced in laboratory. In the wild, selection against hybrids is therefore likely to be an important reproductive barrier between these two morphs.

We reared the offspring of SB-, PL charr and their hybrids under common garden conditions to test for differences between morphs in the values of a wide range of traits, and to assess to what extent these differences could be related to patterns of trait covariance. At the same time, we characterised the trait values of hybrids relative to each morph and assessed whether and how disrupted patterns of trait covariance may affect their development. Intermediate or transgressive individual traits can be studied as average trait values [56]. However, because developmental disturbances can result in increased individual deviations from group means [57], we also tested whether hybrids differ from the parent morphs in their trait variances. This information on trait-level average values, trait-level variances and trait covariances should provide an integrative view on the state of phenotypic divergence between the two morphs and on the plausibility of maladapted hybrid phenotypes.

While keeping track of every individual within the common-garden setup, both along the embryonic development (from hatching to the onset of exogeneous feeding) and during their first months as juveniles (until three months after the onset of exogeneous feeding), we measured traits related to morphology and development (hatching date, initial size and growth, yolk sac size and resorption, developmental trajectory of the head shape). These measurements enabled us not only to test for differences in average value, variances and covariances of traits between types of crosses, but also to assess whether and how these traits covary with other traits measured later in life, and which were related to morphology (shape of the feeding apparatus), behaviour (feeding intensity) and growth after the onset of exogeneous feeding. These individual-level measurements were interpreted in a single framework by building morph- or hybrid-specific phenotypic (**P**) matrices of variance-covariance with components referring to traits from different developmental stages.

Firstly, we hypothesized that the two morphs would show diverged patterns of trait covariance reflecting adaptations to the pelagic and the benthic habitats. To the contrary, the observation of conserved **P** matrices in this recent but highly diverged system would suggest that phenotypic constraints can extensively persist along the processes of adaptive divergence. Secondly, we hypothesized that the F_1_ hybrids would show relaxed trait covariances according to the general prediction of genetic architecture breakdowns [9]. However, the hybrids may also show conserved or even novel phenotypic constraints, which in such cases could shed light on developmental deficiencies (e.g. integrated transgressive traits, extreme characters in embryos with consequences on the phenotype at later life stages).

## Methods

### 1. Study system

Thingvallavatn is a deep postglacial lake (surface 84km^2^, mean depth: 34 meters) that formed within a graben of the Mid-Atlantic ridge during the last glacial retreat (*ca*. 10,000 years BP) [58,59]. The lake is characterized by a wide pelagic zone and three major benthic habitats: a “stony littoral” zone (0-10m deep) composed of a spatially complex lava substrate with loose stones, crevasses and interstitial spaces, a deeper zone (10-20m deep), densely vegetated by the algae *Nitella opaca*, and a profundal zone (25 m and deeper) covered by a diatomic gyttja substrate [50,60]. The lake hosts four morphs of Arctic charr. Two of them, the planktivorous (PL) and the piscivorous charr (PI) feed in the pelagic and epibenthic layers, respectively, and are characterised by a terminal mouth and relatively small pectoral fins [61]. The two other morphs, the large-benthic (LB) and the small-benthic charr (SB), forage in the benthic zone, and show a blunt snout with a subterminal mouth and large pectoral fins [48– 50]. The PL and the SB charr are currently found exclusively in sympatry, although coalescent simulations supports evolutionary scenarios involving short periods of geographic isolation [45]. The differentiation of the craniofacial morphology among the two morphs is initiated early during development, before hatching [52], but can also be influenced to some extent by plasticity after the onset of exogeneous feeding [62]. The SB charr spawn from August to November and the PL charr from September to October [55]. The young of the year of the two morphs are believed to use the same habitat, the surf zone (0-1m deep), from the onset of active feeding in the spring until the summer, when the PL-charr are thought to migrate towards the pelagic and the epibenthic zones [63].

### 2. Fish collection and rearing

We collected mature SB and PL charr with gillnets during five sessions of night fishing in October 2017, at a single spawning site known to be used by both morphs (Svínanesvík, 64°11’24.6”N; 21°05’40.5”W; [55]). We used 52 fish to generate 26 full-sib families on site (crossing design in Table S1). The eggs were kept in a vertical incubator (MariSource, USA) and maintained at 4.1±0.2°C at the rearing facility of Hólar University (Verið), Sauðárkrókur, Iceland. On the mean hatching day (when 50% of the embryos from a given family had hatched), 40 free-swimming embryos from each one of the first nine families to hatch were moved into single-individual cylinders with a plastic mesh on the lower side to allow water flows (2.2 cm diameter x 6.0 cm height, 0.1 cm^2^ mesh size), and placed into a EWOS tray (60 x 250 cm) with flow-through water. Before first feeding (ca. 530 degree days– °C *d*, March 2018), embryos were moved into 22cl transparent plastic cups placed in the same EWOS tray (6.1±0.6 °C). These cups were perforated on the sides and were assumed to enable the exchange of olfactory cues and visual contact between congeners. The fish were fed *ad libitum* two or three times a day with ground aquaculture pellets (Inicio Plus G 0.4mm, BIOMAR).

### 3. Data collection

We collected multivariate longitudinal individual-based data on ontogeny (standard length, yolk sac resorption, growth before and after the onset of exogeneous feeding, timing of the onset of exogeneous feeding), trophic morphology (head shape) and feeding behaviour (feeding activity and feeding performance).

We measured the craniofacial development, pre- and post-feeding growth, and yolk-sac resorption using morphometric data from photographs taken at four points throughout ontogeny: at hatching (ca. 445 °C *d*), 20 days post-hatching (ca. 530 °C *d*), three to four weeks after the onset of exogeneous feeding (ca. 840 °C *d*) and two months later (ca. 1100 °C *d*). The fish were anaesthetized with 2-phenoxyethanol [1], positioned on their lateral side facing left and photographed with a fixed, down-facing camera (Canon EOS 650D + 100mm macro lens) before being returned to their respective growing cell. To correct for the tilt caused by the yolk-sac, the specimens were positioned on 3% methyl cellulose [64] for the photographs of the first two timepoints.

The timing of the onset of exogeneous feeding was determined through “One-zero” sampling (*i*.*e*. records of the occurrence or non-occurrence of an event within defined observation periods) [65]. Direct observations were made every day on all fish, starting when food was introduced in the rearing setup for the first time (ca. 635 °C *d*). This was done in the following way: a three minute observation trial was initiated on each focal individual as the observer introduced food (ca. 10 slowly sinking ground pellets particles of 0.4 mm or less) into the cup of the focal fish. We determined the date of the onset of exogeneous feeding as the date the focal fish was observed catching food for the first time.

Several key aspects of feeding behaviour were estimated by conducting three focal sampling sessions [65] over three consecutive days, seven days after the date of first feeding of the focal individual. A three minute observation period was initiated following the introduction of the food, to record the time it took the fish to seize the first particle (reaction time) [26]. From this point on, an extra one-minute observation trial was initiated, during which feeding intensity (number of particles caught) and feeding strategy (proportion of particles caught on the bottom, on the surface and in mid-water) were recorded. The focal fish was considered “nonfeeding” and the trial was terminated if no particle was seized by the end of the initial three-minute observation period.

### 4. Digitizing and pre-processing morphological data

Data on size (standard lengths) and morphology were extracted from photographs using Geometric morphometrics methods [66]. We digitized the contours of the eye, of the head (from the lower edge of the maxilla below the centre of the eye to the point of maximum curvature between the brain and the cranium) and of the yolk sac (from the junction with the vitellin vein to posterior junction with the body) with Bezier curves using the R package Stereomorph. Two landmarks were placed on the extremity of the notochord and on the anus (Figure S2). During the standard pre-processing steps (*i*.*e*. superimposing the landmark configurations of all specimens to a common coordinate system through Generalized Procrustes Analysis) [67], we estimated the surface of the yolk sac as the area of a polygon composed of 200 semi-landmarks extracted from its respective curve. We calculated the standard length of all specimens as the Euclidian distance between the extremity of the notochord and the furthest of 50 semi-landmarks generated from the curve along the head.

The information on shape of the head shortly after the onset of exogeneous feeding was extracted from the same specimens for the global analysis of trait covariance. We generated variables of local shape deformations compared to the grand mean using a set of 16 and 4 semi-landmarks extracted from the curves of the head and the eye, respectively, and corresponding to the third sampling time point (three to four weeks after the onset of exogeneous feeding) [68]. Local shape changes were estimated with spatial interpolation, based on the thin-plate Splines interpolation function developed in LORY [69]. The resulting multivariate dataset was composed of 24 Jacobian determinants (*i*.*e*. scalars describing local shape expansions/contractions) enabling the quantification of explicit, local shape deformation indices (unlike Procrustes shape variables [69]). The matrix of Jacobian determinants was then reduced with Factor Analysis, relying on oblique rotations (Oblimin) and on the Weighted Least Square algorithm, using the R package psych [70]. This procedure produced three independently interpretable variables (WLS1, WLS2 and WLS3) describing three groups of areas of covarying shape changes (Figure S3). The latent variable WLS1 described two areas that were negatively correlated in their extensions/contractions and which corresponded to the cranium and to the maxilla, respectively. The variable WLS2 described extensions/contractions of the frontal region that covary negatively with the extensions/contractions of the nasal region. Finally, WLS3 corresponded to extensions/contractions of an area including the eye.

### 5. Analyses of individual traits

We modelled the growth trajectories of every specimen in each type of cross using polynomial random regressions [71]. We then tested for overall differences between cross type in the development of the head by conducting phenotypic trajectory analyses of the Procrustes residuals of the head [67]. Morphological disparity analyses [72] were used to compare the types of crosses on the basis of within-group variations in head shapes at the third developmental time-point (soon after first feeding). Differences in local deformations in the head were also analysed with three separate generalized linear mixed effect models (GLMM) [73], each containing one of the three latent variables of Jacobian determinants as a response (WLS1, 2 and 3).

We also tested for group differences in the date of first feeding, feeding intensity, and foraging behaviour with separate GLMMs. The specifications of each model are described in Table S3.

All the GLMMs were run with the R package MCMCglmm [74]. MCMCglmm relies on a Bayesian framework using Markov chain Monte Carlo (MCMC) methods. We always set weakly informative priors (*V* = 1, *nu* = 0.002 or the number of traits for the multi-response models) and determined the optimal number of iterations for model convergence through the examination of trace plots, posterior density plots and effective sample sizes (Table S3). Inferences were made by comparing the posterior mode estimates and 95% Highest Posterior Density Credible intervals (95% CrI) of each cross type (and in relation to the zero baseline for the significance of *R* estimates).

We studied between-individual variations in feeding behaviour by comparing repeatability estimates among the three cross types. The repeatability of each behavioural variable measured across the three repeated observational trials (propensity to start feeding, number of caught items, vertical location) was calculated according to the formula of adjusted repeatability in [75]. The repeatability estimates of the propensity to start feeding, a variable with binary data, were calculated accounting for Jensen’s inequality when transforming the results (initially on the latent scale) to the data scale, following [76].

### 6. Trait covariance

We studied the patterns of trait covariance by generating a phenotypic matrix of variance-covariance (**P** matrix) for each cross type. **P** matrices are reliable surrogates of genetically based patterns of trait covariances (*i*.*e*. of the **G** matrices) when no pedigree is available [77,78]. **P** matrices are especially likely to be good proxies in our particular study because the effects of the environment were mitigated by the use of common-garden conditions, and because the parental effects were accounted for by including in the subsequent models the family of origin (*i*.*e*. the egg clutch) of all individuals. We estimated the components of the three matrices by running three separate Multi-Response Generalized Mixed models [74]. All three models contained ten variables as a response (Table 1). The family was included as a fixed effect while the identity of the individual was included as a random factor. All the traits were mean-standardized by dividing the raw values by their group means [79].

**Table 1.** Variables selected for generating phenotypic variance-covariance matrices (one per cross types).

| Category | Variable name |
| --- | --- |
| Development | Size at hatching (D1) |
|  | Size at the onset of first feeding (D3) |
|  | Growth from hatching (D1) to 20 days post-hatching (D2) |
|  | Growth from the onset of exogeneous feeding (D3) to two months after the onset of exogeneous feeding (D4) |
|  | Yolk sac size at hatching (D1) |
|  | Yolk sac conversion |
| Morphology | WLS1: Extensions/contractions of areas in the cranium |
|  | WLS2: Extensions/contractions in the frontal and nasal regions |
|  | WLS3: Extensions/contractions of areas of the eye. |
| Behaviour | Feeding intensity: Number of food items collected during observational trials. |

The **P** matrices of each type of cross were first compared on the basis of their size, shape and orientation [80]. The matrices sizes (*V*_*tot*_) were used to compare the types of crosses in the overall phenotypic variance and were calculated as the sum of their eigenvalues (Equation 2 in [81]) [80,81]. Eccentricity (*Ω*) was used as a measure of the shape of the matrices and was calculated as the ratio of their first eigenvalue over the sum of the eigenvalues (Equation 3 in [81]). Differences in overall matrix orientation were assessed using the angles (*θ*) between the first eigenvector of each **P** matrix. Briefly, if the patterns of trait covariances were not conserved but have rapidly evolved among the two morphs, we expected the two types of pure-morph offspring to show differences in the overall size of **P** (*V*_*tot*_), which should suggest a response to two selective regimes eroding genetic variations to different extents. Similarly, differences in eccentricity (*Ω*) between the two pure-bred offspring were expected (for example, correlational selection, which can produce more constrained, “cigar shaped”, **G** matrices [80], might differ among the respective habitats of each morph). The orientation of **G** can also be subjected to changes because of the effects of correlational selection, among other evolutionary forces [68,80,82]. Thus, differences between pure-bred offspring in the orientation of **P** (*θ*) were also expected [83]. Regarding the hybrids, breakdowns in their trait covariance structure should be indicated by **P** matrices with larger sizes and reduced eccentricity [35]. Meanwhile, differences in the orientation of **P** between the hybrid and the pure-morph offspring should indicate whether the remaining constraints on the hybrid phenotypes are intermediate, under dominance and conserved relative to one morph, or transgressive (*i*.*e*. biased toward a unique direction of the phenotypic space).

We then tested for among group (the two pure morph progeny and the hybrids) dissimilarities in particular components of the **P**-matrices using the Random skewers method, an approach enabling the identification of differences in the variance-covariance structure of matrices that may reflect differences in phenotypic constraints and putative responses to selection [84]. The Random skewers method consists in projecting random vectors of selection through **P** or **G** to uncover specific regions of the (phenotypic) space where matrices differ in their variances and covariances. Here we applied the Random skewers approach within a Bayesian framework, following [85]. Each one of 1000 random vectors were projected through each one of the MCMC 1000 estimates of **P** for the three types of cross. The resulting vectors that did not overlap in their 95% Credible intervals between any pair of type of crosses were then collated and their variance-covariance calculated to generate the *n* x *n* **R** matrix (*n* = number of traits). This **R** matrix describes the parts of the phenotypic space where populations differ in variance-covariance for specific trait combinations.

For visualisation purpose, **P** matrices were projected into a subspace composed by the first three eigenvectors **P** matrix of the PLxPL offspring by modifying the plotsubspace() function from [74]. Because angles between eigenvectors are necessarily positive, we also estimated angles between the estimated **P** matrices and **P** matrices of simulated populations containing 150 individuals from each pair of cross types whose angle was studied (code available in the supplementary material).

## Results

### 1. Differences at the level of individual traits

The SBxSB and the hybrid offspring significantly differed from the PLxPL offspring in their growth trajectories (Figure 2, details of the regression estimates in Table S4). The SBxSB and hybrid offspring tended to have lower intercepts than PLxPL offspring (posterior modes [95% CrIs] of log_10_(standard length) = PLxPL: 3.09 [2.97; 3.19], SBxSB: 2.98 [2.90; 3.11], hybrids: 2.98 [2.92; 3.07]). Furthermore, lower slopes and small second order polynomial terms were observed in the SBxSB and the hybrid offspring compared to the PLxPL offspring (slopes = PLxPL: 6.14 [5.89; 6.38], SBxSB: 5.77 [5.38, 5.95], hybrids: 5.73 [5.55; 5.92]; second order polynomial terms = PLxPL: −0.70 [-0.85; −0.55], SBxSB = −1.14 [-1.37; −0.99], hybrids: −1.00 [-1.11; −0.85]). These results indicate a slower and more decelerating growth in the SBxSB and the hybrid offspring than in the PLxPL offspring.

**Figure 2.**
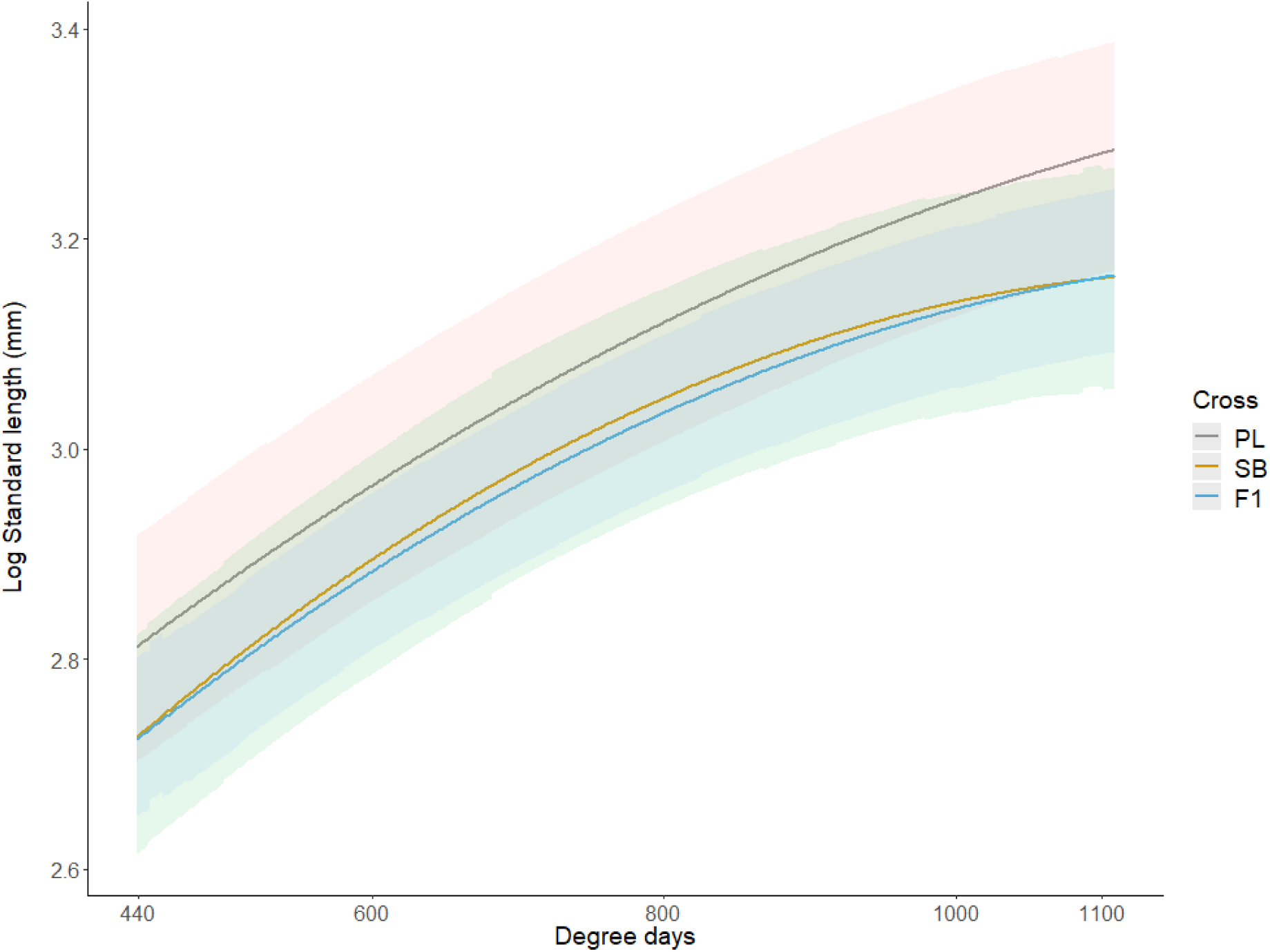
Growth trajectories of each cross type (predicted values and 95% Confidence intervals). The growth period under study started at hatching (ca. 400 degree days) and ended three months after the onset of exogeneous feeding (ca. 1100 degree days). PL: PLxPL offspring, SB: SBxSB offspring, F_1_ : F_1_ hybrid offspring.

We did not observe strong differences between the types of crosses in mean yolk sac area at hatching nor in the resorption (Table 2). The hybrids and the SBxSB offspring appeared to have smaller yolk sac sizes at hatching and the hybrids tended to have faster resorption rate, although wide overlaps in 95% intervals confer low levels of certainty to these patterns.

**Table 2.**
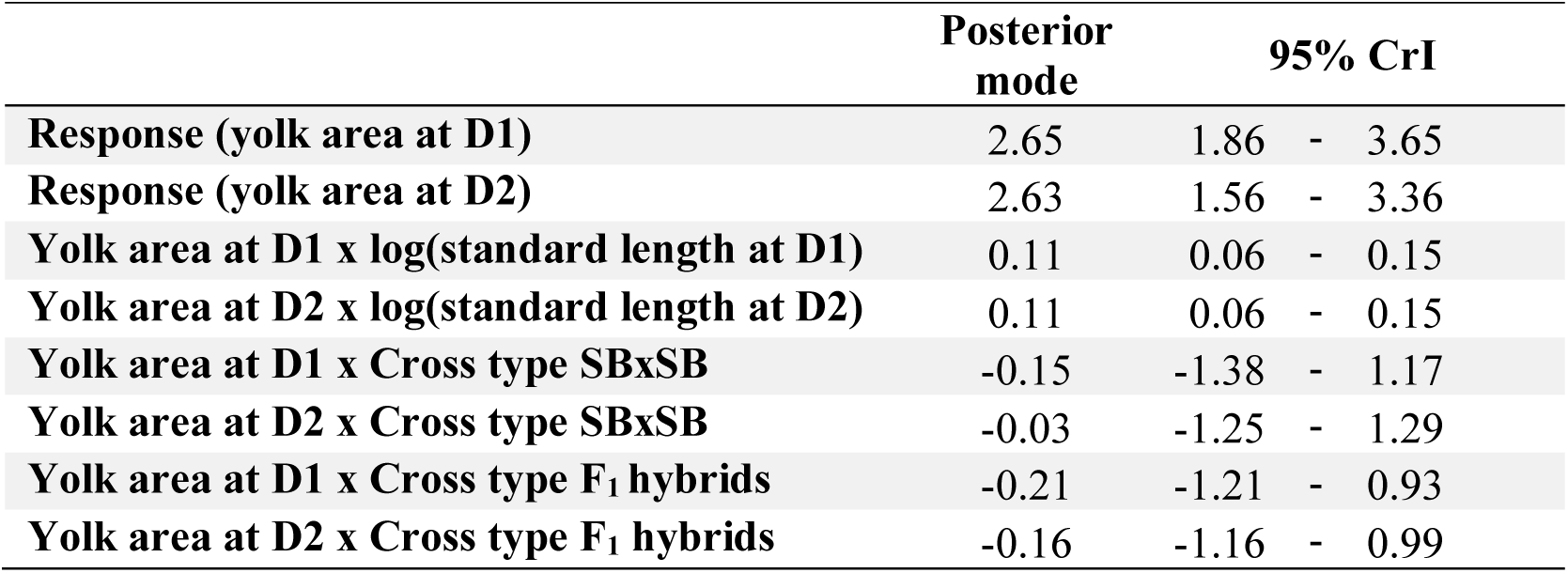
Posterior estimates of the fixed effects from the Multi-response Generalized Linear Mixed Effect Model on the yolk sac area (mm^2^). The PLxPL cross type is the base line. D1: hatching, D2: 20 days post-hatching. See Table S3 for the details of the model.

|  | Posterior<br>mode | 95% CrI |  |  |
| --- | --- | --- | --- | --- |
| Response (yolk area at D1) | 2.65 | 1.86 | - | 3.65 |
| Response (yolk area at D2) | 2.63 | 1.56 | - | 3.36 |
| Yolk area at D1 x log(standard length at D1) | 0.11 | 0.06 | - | 0.15 |
| Yolk area at D2 x log(standard length at D2) | 0.11 | 0.06 | - | 0.15 |
| Yolk area at D1 x Cross type SBxSB | -0.15 | -1.38 | - | 1.17 |
| Yolk area at D2 x Cross type SBxSB | -0.03 | -1.25 | - | 1.29 |
| Yolk area at D1 x Cross type F<sub>1</sub> hybrids | -0.21 | -1.21 | - | 0.93 |
| Yolk area at D2 x Cross type F<sub>1</sub> hybrids | -0.16 | -1.16 | - | 0.99 |

Analyses of the Procrustes residuals (Randomized Residuals Permutation Procedure) of the head shapes indicated that size was related to most of the variation among specimens while no effect of the type of cross in itself was observed (Table 3). The ontogenetic trajectories of the head shape did not differ significantly between the types of crosses in shape, path length and angle attributes (Figure 3a, Table S5). No differences between types of crosses in the variances of the head shapes were observed from the disparity analyses at hatching and at the onset of exogeneous feeding (absolute differences in Procrustes variances < 0001, *p*-value > 0.1 in all the pairwise comparisons, Table S6). We did not observe differences among cross types in mean local shape changes based on the GLMMs of the latent variables of correlated Jacobian determinants (Table S7). The same analyses, however, revealed reduced residual variances in the SBxSB offspring compared to the PLxPL offspring and the hybrid offspring (Figure 3, comparisons of the other latent variables in Figure S4). This pattern is supported by limited overlaps in the 95% CrI and indicates lower variations in the extensions/contractions of areas of the eyes in SBxSB offspring compared to the two other types of crosses (Figure 3b).

**Table 3.** Formula and results of the regression on Procrustes residuals of the head shapes in the specimens reared individually. Families are nested within cross type. Age: Sampling time point, Size: Centroid size.

| Formula |  |  |  |  |  |
| --- | --- | --- | --- | --- | --- |
| Procrustes coordinates ~ log<sub>10</sub>(Size) + Cross/Family + log<sub>10</sub>(Size) x Cross/Family + Age x Coss/Family |  |  |  |  |  |
| Table of variance |  |  |  |  |  |
| Effects | d.f. | SS | R<sup>2</sup> | Z | p |
| log(Size) | 1 | 5.17 | 0.62 | 7.80 | <0.01 |
| Cross type | 2 | 0.06 | 0.01 | -1.18 | 0.88 |
| Age | 3 | 0.63 | 0.07 | 9.47 | <0.01 |
| Cross type x Family | 5 | 0.25 | 0.03 | 8.07 | <0.01 |
| Cross type x log(Size) | 2 | 0.02 | 0.00 | -2.64 | 1.00 |
| Cross type x Age | 6 | 0.07 | 0.01 | -3.09 | 1.00 |
| Cross type x log(Size) x Family | 5 | 0.06 | 0.01 | 4.08 | <0.01 |
| Cross type x Age x Family | 15 | 0.10 | 0.01 | 3.63 | <0.01 |
| Residuals | 626 | 2.06 | 0.24 | - | - |
| Total | 665 | 8.40 | - | - | - |

**Figure 3.**
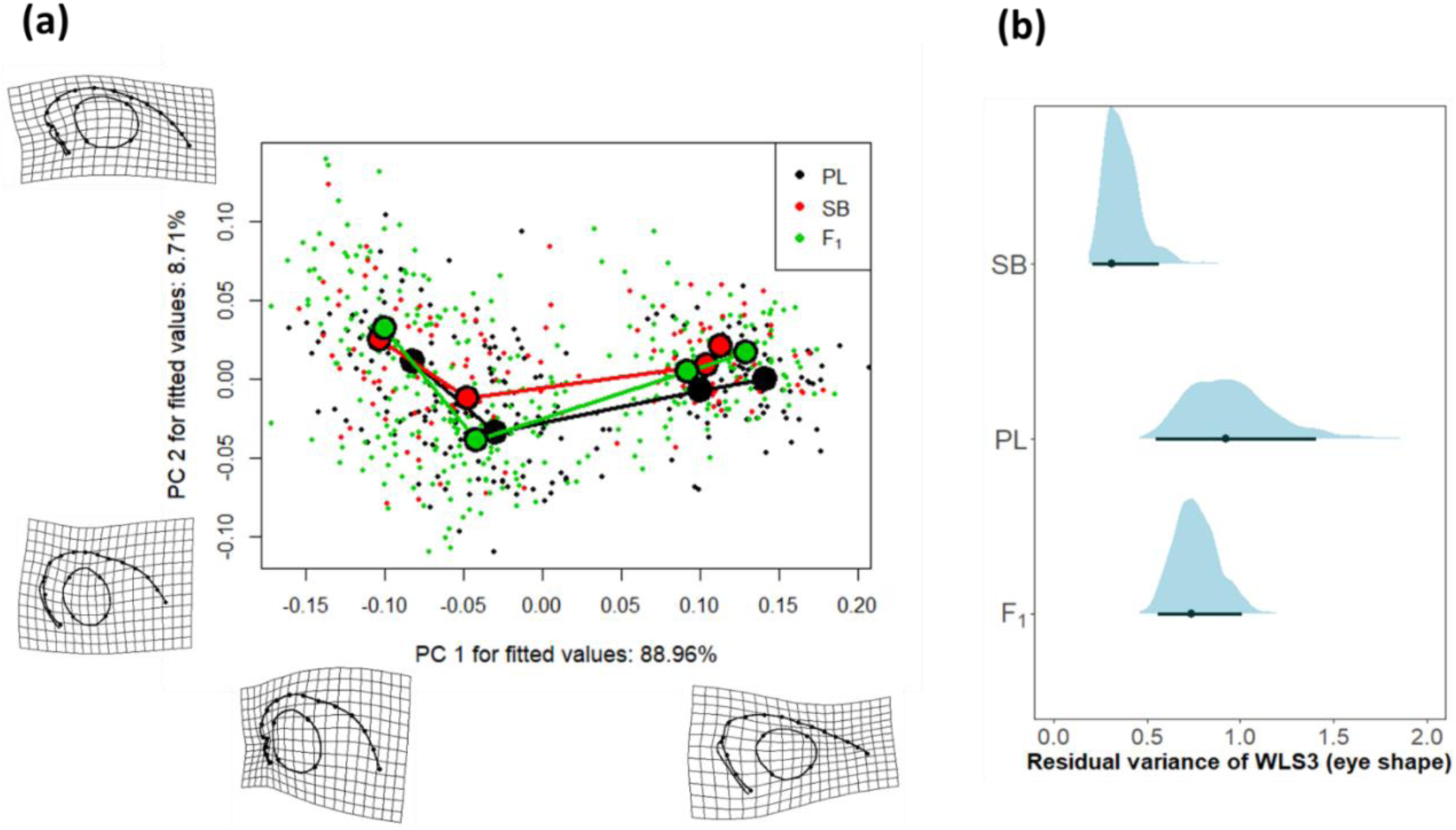
Variation in head shape among the three cross types. (a) Principal component (PC) plot of the ontogenetic trajectories across the four time points and deformation grids of the extremes of each PC axis (time points from left to right: hatching, 20 days post hatching, onset of exogeneous feeding, two months after the onset of exogeneous feeding). Because size variations among morphs are of interest, the shapes are not corrected for size. (b) Posterior densities, posterior mode and 95% credible interval of the residual variance of WLS3 (contractions/extensions in areas of the eye). Categories: SB = SBxSB offspring, PL = PLxPL offspring, F_1_ = hybrid offspring.

We did not observe differences among cross types either in the mean or in the variances of the date of the onset of exogeneous feeding, the estimated date being very close to one another (Table S8). There was no apparent difference between groups in the propensity to start feeding during the experimental trials on feeding behaviour (Figure S5). However, the PLxPL offspring showed a higher level of consistent individual differences (repeatability) in their propensity to start feeding (R = 0.41 [0.23; 0.53], posterior mode [95% CrI]) than the fish of the SBxSB offspring (R = 0.00 [0.00; 0.25]) and the hybrids (R = 0.00 [0.00; 0.27]). The estimated number of captured food items also appeared lower for the fish of PLxPL offspring than for their SBxSB conspecifics (Figure 4a). The overlap in the 95% CrI of the two types of pure-morph offspring provides moderate statistical support for these differences, while no strong inference can be made from the apparently intermediate level of feeding intensity in hybrids (Figure 4a). PLxPL individuals also tended to show lower variance than the SBxSB individuals and the hybrids in the number of attacked items (Figure 4b). The comparison of these estimates indicated that the three types of crosses showed low to null levels of consistent differences between individuals in this feeding behaviour (Figure 4c). We did not observe differences between cross types in the propensity to use the bottom of the container, the water column or the surface of the water when foraging (Figure S5a-i).

**Figure 4.**
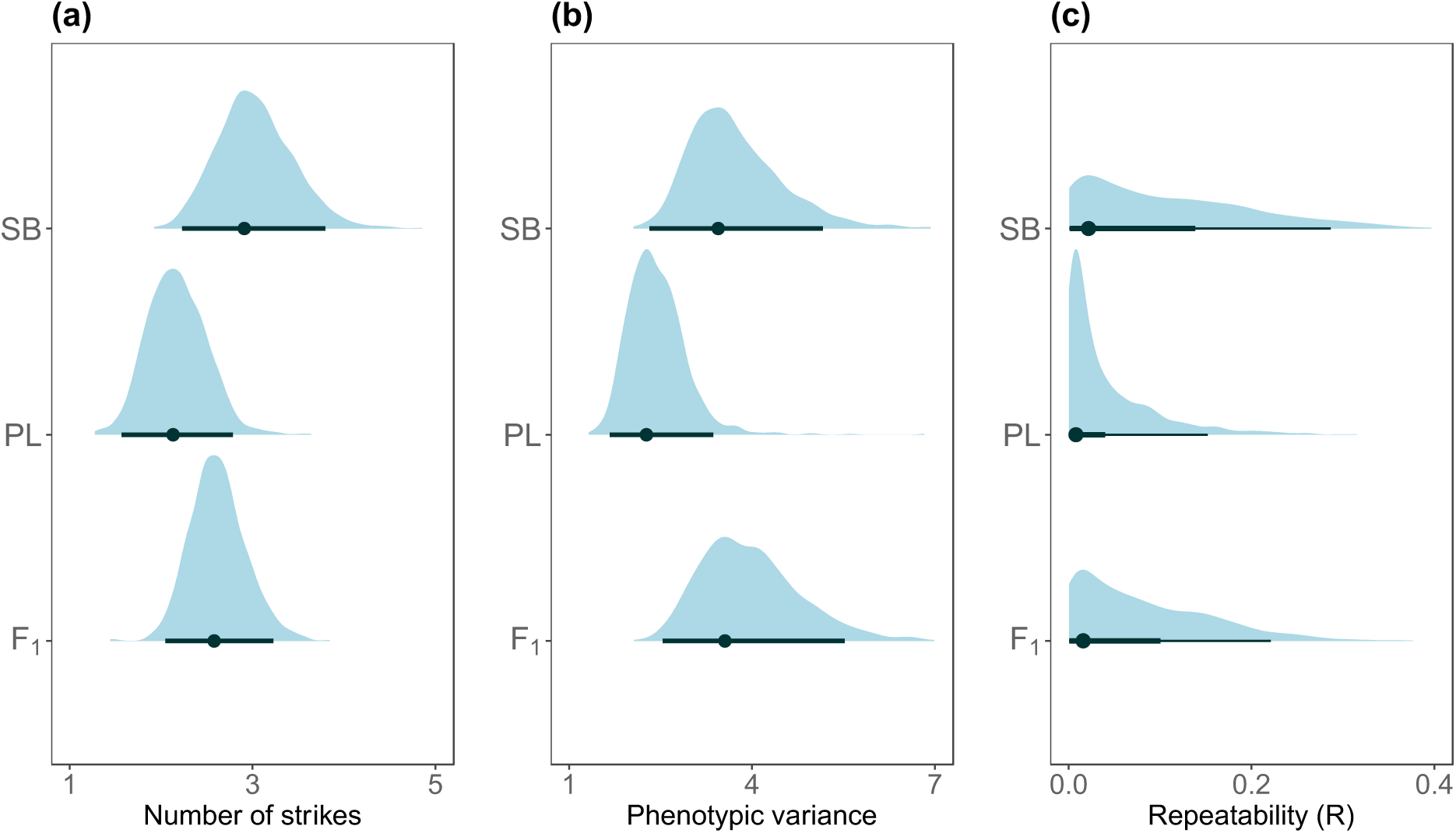
Posterior densities, posterior modes and 95% CrIs of the number of food items attacked by the focal fish during the feeding trial and among the three types of crosses. (a) Number of attacks, (b) total variance, and (c) Repeatability (R) as a measure of consistent between-individual differences. Categories: SB = SBxSB offspring, PL = PLxPL offspring, F_1_ = hybrid offspring.

### 1. Trait covariance in pure-morphs crosses and hybrids

We observed differences among cross types in variance regarding body size and growth (before and after the onset of exogenous feeding), yolk sac area at hatching and yolk sac conversion (Table 4). Higher variance in growth during exogeneous feeding was observed in the hybrids. However, the hybrids were intermediate in variance or non-different from at least one morph regarding the other traits.

**Table 4.**
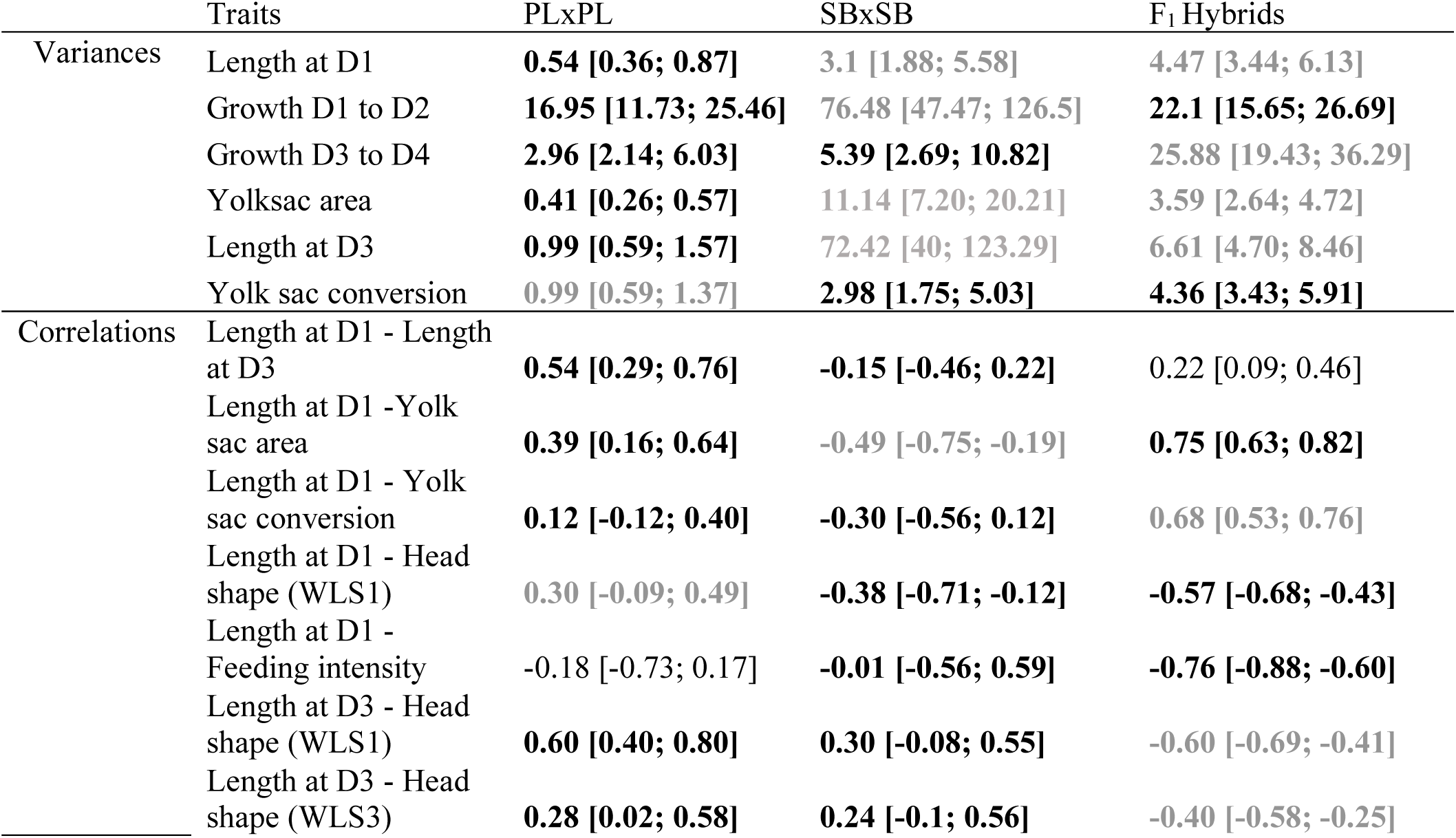
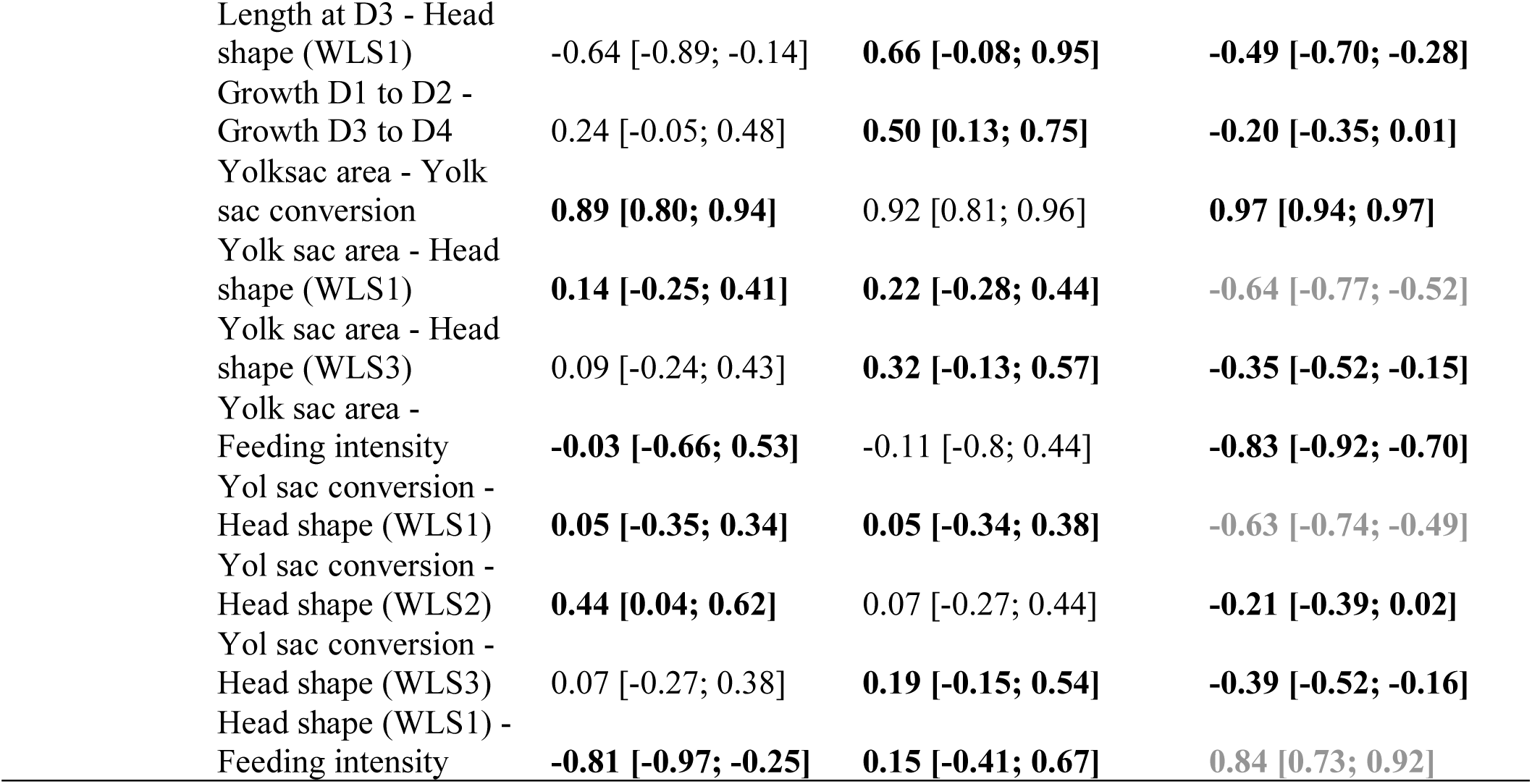
Posterior modes and credible intervals (CrIs) of trait variance and correlations that showed nonoverlaps in 95% CrIs between at least two cross types. Developmental time points D1 = hatching, D2 = 20 days post hatching, D3 = onset of exogeneous feeding, D4 = two months after the onset of exogeneous feeding. WLS1: Extensions/contractions of areas of the cranium. WLS3: Extensions/contractions of areas in the eye. WLS2: Extensions/contractions in the frontal and the nasal region. Estimates with non-overlapping 95%CrIs are in bold and grouped by shades of grey.

|  | Traits | PLxPL | SBxSB | F<sub>1</sub> Hybrids |
| --- | --- | --- | --- | --- |
| Variances | Length at D1 | **0.54 [0.36; 0.87]** | 3.1 [1.88; 5.58] | 4.47 [3.44; 6.13] |
|  | Growth D1 to D2 | **16.95 [11.73; 25.46]** | 76.48 [47.47; 126.5] | **22.1 [15.65; 26.69]** |
|  | Growth D3 to D4 | **2.96 [2.14; 6.03]** | **5.39 [2.69; 10.82]** | 25.88 [19.43; 36.29] |
|  | Yolksac area | **0.41 [0.26; 0.57]** | 11.14 [7.20; 20.21] | 3.59 [2.64; 4.72] |
|  | Length at D3 | **0.99 [0.59; 1.57]** | 72.42 [40; 123.29] | 6.61 [4.70; 8.46] |
|  | Yolk sac conversion | **0.99 [0.59; 1.37]** | **2.98 [1.75; 5.03]** | **4.36 [3.43; 5.91]** |
| Correlations | Length at D1 - Length at D3 | **0.54 [0.29; 0.76]** | -0.15 [-0.46; 0.22] | 0.22 [0.09; 0.46] |
|  | Length at D1 - Yolk sac area | **0.39 [0.16; 0.64]** | -0.49 [-0.75; -0.19] | **0.75 [0.63; 0.82]** |
|  | Length at D1 - Yolk sac conversion | **0.12 [-0.12; 0.40]** | -0.30 [-0.56; 0.12] | **0.68 [0.53; 0.76]** |
|  | Length at D1 - Head shape (WLS1) | **0.30 [-0.09; 0.49]** | -0.38 [-0.71; -0.12] | **-0.57 [-0.68; -0.43]** |
|  | Length at D1 - Feeding intensity | -0.18 [-0.73; 0.17] | **-0.01 [-0.56; 0.59]** | **-0.76 [-0.88; -0.60]** |
|  | Length at D3 - Head shape (WLS1) | **0.60 [0.40; 0.80]** | **0.30 [-0.08; 0.55]** | **-0.60 [-0.69; -0.41]** |
|  | Length at D3 - Head shape (WLS3) | **0.28 [0.02; 0.58]** | **0.24 [-0.1; 0.56]** | **-0.40 [-0.58; -0.25]** |

|  |  |  |  |
| --- | --- | --- | --- |
| Length at D3 - Head shape (WLS1) | -0.64 [-0.89; -0.14] | **0.66 [-0.08; 0.95]** | **-0.49 [-0.70; -0.28]** |
| Growth D1 to D2 - Growth D3 to D4 | 0.24 [-0.05; 0.48] | **0.50 [0.13; 0.75]** | **-0.20 [-0.35; 0.01]** |
| Yolksac area - Yolk sac conversion | **0.89 [0.80; 0.94]** | 0.92 [0.81; 0.96] | **0.97 [0.94; 0.97]** |
| Yolk sac area - Head shape (WLS1) | **0.14 [-0.25; 0.41]** | **0.22 [-0.28; 0.44]** | **-0.64 [-0.77; -0.52]** |
| Yolk sac area - Head shape (WLS3) | 0.09 [-0.24; 0.43] | **0.32 [-0.13; 0.57]** | **-0.35 [-0.52; -0.15]** |
| Yolk sac area - Feeding intensity | **-0.03 [-0.66; 0.53]** | -0.11 [-0.8; 0.44] | **-0.83 [-0.92; -0.70]** |
| Yol sac conversion - Head shape (WLS1) | **0.05 [-0.35; 0.34]** | **0.05 [-0.34; 0.38]** | **-0.63 [-0.74; -0.49]** |
| Yol sac conversion - Head shape (WLS2) | **0.44 [0.04; 0.62]** | 0.07 [-0.27; 0.44] | **-0.21 [-0.39; 0.02]** |
| Yol sac conversion - Head shape (WLS3) | 0.07 [-0.27; 0.38] | **0.19 [-0.15; 0.54]** | **-0.39 [-0.52; -0.16]** |
| Head shape (WLS1) - Feeding intensity | **-0.81 [-0.97; -0.25]** | **0.15 [-0.41; 0.67]** | **0.84 [0.73; 0.92]** |

The two pure-morph offspring differed in four off-diagonal elements of **P**, which corresponded to correlations between body sizes before and after exogeneous feeding, body size and yolk sac area at hatching, and body size at hatching and head morphology (Table 4). However, sixteen off-diagonal elements of **P (***i*.*e*. correlations) differed between the hybrids and the offspring of at least one of the pure-morph crosses. These elements corresponded to correlations involving body size at hatching, growth, head shape, feeding intensity, size and yolk sac conversion. Four of these elements described stronger correlations between the hybrids and both morphs. These were correlations between body size at hatching and yolk sac conversion, yolk sac area and head shape, and head shape and feeding intensity. Trends regarding all the other trait correlations can be visualized in Figure S6.

We also observed differences in the summary estimates of **P** (size, eccentricity and orientation) among cross types. First, we observed a higher estimate of *V*_*tot*_ (matrix size) in the SBxSB offspring compared to the PLxPL, while the hybrids appear to be intermediate for this metric (Figures 5a-c and 5d). This indicates higher phenotypic variance overall in the SBxSB than in the PLxPL and the hybrid offspring. Second, eccentricity (*Ω*) was lower in the hybrids than in the PLxPL offspring (Figures 5a-c and 5e) and tended to be lower than in the SBxSB offspring (overlap in 95% CrI provided modest support to this trend, Figure 5e). This pattern suggests that a small number of traits drive phenotypic variation to a lesser extent in the hybrids than in the PLxPL offspring. However, the three cross types did not differ in the orientation of **P**, the angles among of the first eigenvector of each pair of cross type (*θ*) being low and nondifferent from the angles between the observed and the random **P** matrices (Figure 5f).

**Figure 5.**
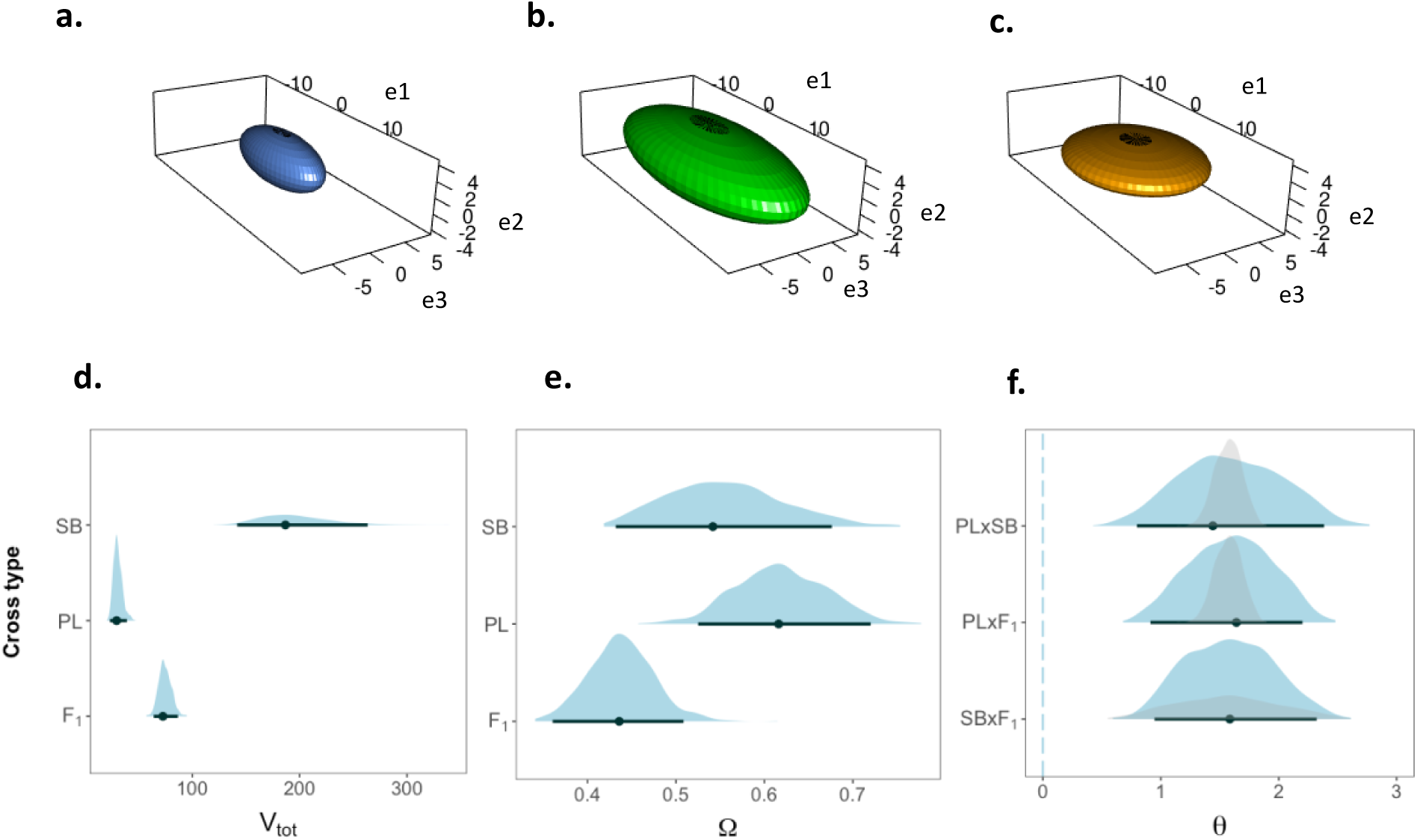
Summary property the **P** matrices of each type of cross. (a-c) Ellipsoid representations of the posterior modes of each matrix projected in a subspace defined by the first three eigenvectors of **P** from the PLxPL cross. The axes explain 83%, 100% and 100% of the variance of **P** in the PLxPL, the SBxSB and the hybrid crosses, respectively. The eigenvectors are detailed in Figure S7. (d-f) Posterior densities, posterior modes and 95% CrIs of the three summary estimates of the matrices of phenotypic variance of each cross type, being (d) the overall phenotypic variance (*V*_*tot*_), (e) the eccentricity *(Ω*), and (f) the angle (*θ*) between the first eigenvectors. Densities of the angle estimates between **P** of a cross type (from top to bottom. PLxPL, PLxPL, SBxSB) and a random matrix are shown in light grey. Cross type: SB = SBxSB offspring, PL = PLxPL offspring, F_1_ = hybrid offspring.

At a finer scale, differences in particular components of the variance-covariance structure of **P** were revealed using the Random skewers method. 995 of the 1000 random vectors had non-overlapping 95% CrIs for at least one pairwise comparison of phenotypic variances among the three cross types. The eigenanalysis of the **R** matrix generated with these vectors revealed differences in phenotypic variance between the three cross types. The first three eigenvectors of **R (e1, e2** and **e3)** accounted for 30% of the overall variation. These three vectors mainly described associations between body size at hatching, growth, yolk sac size and hatching and conversion, and head shapes (mostly extensions/contractions of the frontal regions, Table 5). Note, however, that feeding intensity was also an important component of **e1**. Projecting the **P** matrices on **R** revealed a higher phenotypic variance in the SBxSB offspring than in the PLxPL offspring in the direction of the three vectors (Figure 6a-c). Phenotypic variance in the direction of these vectors appeared to be intermediate in the hybrids, although overlaps in 95% CrI between the hybrid and the pure-bred offspring were observed in the direction of **e1** and **e2**.

**Table 5.** Trait loadings of the first two eigenvectors of **R**. Developmental time points D1 = hatching, D2 = 20 days post hatching, D3 = onset of exogeneous feeding, D4 = two months after the onset of exogeneous feeding.

|  | Eigenvector |  |  |
| --- | --- | --- | --- |
|  | e1 | e2 | e3 |
| Variance explained | 0.11 | 0.11 | 0.11 |
| Size at D1 | -0.47 | -0.39 | 0.41 |
| Size at D3 | 0.28 | -0.37 | -0.10 |
| Growth D1 to D2 | -0.07 | -0.29 | 0.13 |
| Growth D3 to D4 | -0.17 | -0.44 | -0.13 |
| Yolk sac size at D1 | -0.38 | 0.14 | 0.00 |
| Yolk sac conversion | -0.12 | -0.48 | -0.61 |
| Head shape (WLS1 : cranium and maxilla) | -0.22 | 0.12 | 0.11 |
| Head shape (WLS2 : frontal and nasal regions) | -0.44 | 0.41 | -0.45 |
| Head shape (WLS3 : eye) | 0.06 | -0.02 | 0.43 |
| Feeding_intensity | -0.52 | -0.05 | 0.14 |

**Figure 6.**
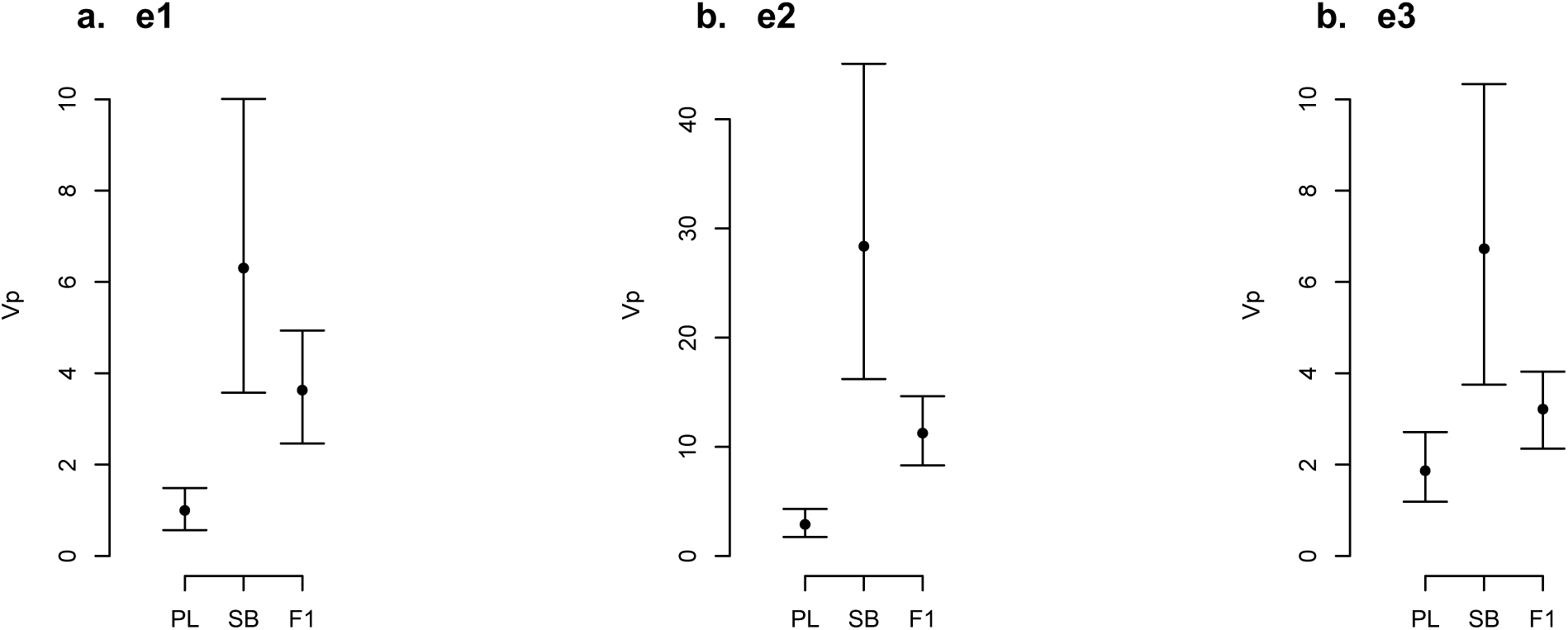
Posterior mean and 95% CrIs of phenotypic variance in the direction of the first (a), the second (b), and the third (c) eigenvector of **R** for the three cross types.

## Discussion

### Traits divergence and hybrid phenotype

Characterizing the developmental origins of hybrid phenotypes is a prerequisite to the understanding how hybridization affects the processes of speciation [56]. Intermediate hybrids from two diverging populations are often seen as disadvantaged compared to the offspring from intra-population mating, but transgressive characters are believed to sometimes facilitate further divergence by enabling the colonisation of new ecological niches [9]. In our common-garden study, the F_1_ hybrids of two sympatric morphs of Artcic charr showed subtle phenotypic differences from the offspring of the two pure morph crosses. First, while SB and PL charr differed in their growth trajectories (which is in line with previous findings on their life-history strategies [49]), the hybrids differed from the PL charr in their growth (although no difference between the hybrids and the SB charr were observed). However, our results did not provide strong evidence for differences in the average values of the yolk sac size and resorption among the three cross types. Subtle differences in head shape were observed, and these were mostly related to lower variations in areas of the eye in the SB charr compared to the PL charr and the hybrids. However, head morphology was dependent on size in the same way for the three cross types (common allometry). The two morphs may therefore differ in shape because of their differences in growth, although studies involving both stained embryos and living juveniles at older ages indicate size-independent differences in the craniofacial morphology of the PL- and SB charr [52,62].

The PL charr also show higher individual consistency in their propensity to start feeding and appeared to be less active and less variable in their feeding behaviour than the SB charr, which corroborates previous observations suggesting that the two morphs have evolved different foraging strategies [54]. Regarding these patterns, the hybrids tended to show intermediate mean values in feeding intensity but appeared to be similar to the SB charr in their variance.

Hints of potentially disadvantaged hybrid phenotypes are therefore already observed at the level of individual traits. These results are in line with other common-garden experiments reporting potentially maladapted hybrids with intermediate or transgressive characters [8], such as intermediate morphologies and lower feeding performances in hybrids from diverging morphs of stickleback fish (*Gasterosteus spp*.) [86], and extreme hatching dates in hybrids from ecotypes of lake whitefish (*Coregonus cluteaformis*) [7]. However, the F_1_ hybrids from our study were not strictly intermediate or transgressive but rather appeared to be similar to one morph or the other, depending on the type of traits, e.g. similar to the SBxSB offspring in their growth, yolk sac resorption and feeding behaviour, but similar to the PLxPL offspring in their morphology. Thus, more complexity in the development of ecologically relevant traits among the two diverging populations and their hybrids may be uncovered by studying a wider range of traits, and a more thorough characterisation of the phenotypes of the three groups could be obtained by adding information on the patterns of covariance among these traits.

Our analyses of the variance-covariance of a large diversity of traits shed more light on those aspects and revealed further phenotypic differences among the two morphs and their hybrids. While the two morphs showed differences in separate trait correlations, their hybrids were either intermediate or similar to one morph. Furthermore, correlations between several traits were stronger in the hybrids than in the two morphs. The summary statistics of the phenotypic matrix of variance-covariance (**P)** indicated higher overall phenotypic variance in the SBcharr than the PL charr, the hybrids being intermediate. Reduced phenotypic constraints also appeared in the hybrids, at least in comparisons with the PLxPL offspring. Finally. random projections of selection vectors through the **P** indicated more phenotypic variations in the SBxSB offspring in the direction of a combination of traits related to growth, yolk sac resorption head morphology and feeding behaviour, while such values tended to be intermediate in the hybrids.

### Evolutionary implications of the differences in trait covariance

While trait covariances can facilitate evolutionary changes they can also constraint the divergence of populations [87]. Recent empirical studies for example report relatively conserved patterns of trait covariances among geographically separated populations, which potentially limit their divergence [81,83]. Our results support these observations to some extent because the two morphs (which are entirely sympatric but present genetic signatures of reproductive isolation [46,47]) differed in their correlation of growth traits across developmental stages, and overall appeared to have evolved only subtle differences in their phenotypic covariance structure. While some differences in trait correlations were observed between the two morphs, their hybrids sometimes showed correlations falling outside of the range of values of the pure-bred offspring, which could be interpreted as extreme characters, just like transgressive separate traits usually studied in the context of speciation. Trait mismatch in hybrids were already observed as being maladaptive in benthic and limnetic species pairs of stickleback fish where divergent selection targets assemblages of genetically independent morphological character [88]. However, our results represent another possibility for selection against hybrids in a multivariate context, where transgressive phenotypes are composed of sets of covarying traits.

The particular pattern of trait covariance observed in the hybrids in our study contrasts with the view that hybridization should break down the genetic architecture underlying trait covariations, and therefore facilitate adaptive diversification through the colonisation of new environments with novel selection pressure [9,35]. Hybrids of several species of African cichlids (*Astatotilapia* sp.) for example show increased phenotypic variations and reduced phenotypic constraints at the morphological level from the generation F_1_-on [35]. Although we observed reduced phenotypic constraints in hybrids of the PL and SB charr, the levels of phenotypic variances were low compared to the SB charr, and novel correlations appeared, suggesting a more complicated picture. Moreover, our results from the Random skewers approach suggested intermediate phenotypic variations in hybrids in a dimension of the phenotypic space corresponding to a combination of developmental, morphological and behavioural traits.

Under phenotypic constraints, evolutionary changes are expected to be biased in the direction of the phenotypic space with the largest variance [78]. The evolution of the hybrid charr may therefore be biased in the direction of this dimension of the phenotypic space, which may limit the room for manoeuvre allowing the colonization of new niches with novel trait combinations.

The pattern of trait covariance associated with apparently low variability in the hybrids therefore appear unlikely to be beneficial in the wild, and further investigations on the direct fitness consequences of transgressive trait correlations in hybrid should bring new insights to the significance of trait covariance in the evolution of post-zygotic barriers between diverging populations. Little information is available on the nature of selection affecting the traits studied here, and it can only be assumed that the phenotypic values observed in the two morphs reflect adaptions to different fitness optima. However, numerous differentiated regions scattered across the genome of the PL and the SB charr have been identified recently [47], suggesting that a large number of traits may have diverged in response to divergent selection. More precisely, the differences in head and body shape between SB and PL charr were shown to result from divergent selection related to trophic and non-trophic environmental variables [89]. Thus, it may be reasonable to assume that the phenotypic values we observe in these two morphs approximated their respective fitness optima, and that deviation from these values as observed in the hybrids may be detrimental. Furthermore, our results from the Random Skewer analyses indicate that phenotypic variation may be biased in hybrid by a combination of traits mostly related to early growth and development/metabolism (*i*.*e*. yolk sac resorption), and that the strongest differences observed in average traits values between the three cross types concerns the growth trajectories, the hybrids growing as slow as the smaller morph (the SB charr). Growth-related traits are known for often being highly related to fitness [22,90], so these results bring further indications that the variations observed in this study may have important ecological implications.

## Conclusion and future directions

Classical studies on the emergence of reproductive isolation as the ultimate result of adaptive divergence often rely on the logical framework that hybrids that are phenotypically different from the parental types may be poorly adapted to either parental niche [3,8,91]. Applying this view to a context of trait covariance, and considering the early development of traits of different nature (ontogeny, morphology, behaviour,), we observed genetically-based phenotypic variations that would not be revealed by studying the mean value of separate traits only. The outstanding pattern of trait covariance observed in the hybrids from our study might originate from nonexclusive effects of correlational selection, differences in the genetic architecture of the two morphs, or developmental and functional constraints [14,16]. These considerations emphases the need for further studies on the proximate mechanisms of trait covariance in a context of ecological speciation. Disentangling these effects through studies of recently diverged sympatric populations should bring new insights to the processes of speciation.

## Acknowledgement

We are very thankful to Skúli Skúlason for his comments of the manuscript and to Neil Metcalfe for his advices for analysing the data on growth and development. We are also thankful to David Benhaïm and Bjarni K. Kristjánsson for their comments on the experimental design and the conceptual aspects of the study. Field work could be conducted thanks to the help of Zophonías O. Jónsson, the members of the Arctic Charr and Salmonids Group of the University of Iceland, and the owners of the farm of Mjóanes, Jóhann Jónsson and Rósa Jónsdóttir. We thank Kári H. Árnason, Rakel Þorbjörnsdóttir, and Christian Beuvard for the organisation and the maintenance of the experimental setup. We are also thankful to Alia Desclos for her help in rearing the fish.

## Authorship Contributions

Quentin J.B. Horta-Lacueva conceived the study, reared the specimens, collected the data, conducted the analyses and drafted the manuscript. Sigurður S. Snorrason coordinated the field work, produced the embryos and critically revised the manuscript. Michael B. Morrissey provided guidance for the data analyses, contributed to the biological interpretations of the results and reviewed the manuscript. Camille A.L. Leblanc developed the experimental setup, contributed in designing the behavioural experiments and reviewed the manuscript. Kalina H. Kapralova established the crossing design, produced the embryos, organized the logistics of the transfer and the maintenance of the specimens, provided guidance during the experiments and critically revised the manuscript. All authors gave their final approval for publication and agree to be accountable for the work therein.

## Ethical note

Sampling was conducted with the permissions of the owner of the farm of Mjóanes and the Thingvellir National Park commission. Ethics committee approvals for research project are not required by the Icelandic regulation (Act No. 55/2013 on Animal Welfare). The rearing and the experimental work were however conducted in the facilities of Hólar University Aquaculture Research Station, which has an operational license under the Icelandic Aquaculture law (Law No. 71/2018). This law includes clauses of best practices for animal care and experimental work. The fish were killed according to the most careful euthanasia guidelines for salmonid fish [1], and the optimal dosage for anaesthesia on 2-phenoxyethanol was adjusted to the reactions of each individual, following the recommendations of the laboratory facility. Decisions on the sample size and on the design of the common-garden experiment were made to ensure that additional studies could be conducted with data collected on the same specimens.

## Founding

This work was fully funded by the Icelandic Centre of Research, RANNÍS (Icelandic Research Fund grant no.173802-051).

## Data accessibility

The data will be deposited onto the Dryad Digital Repository upon acceptance.

