## supplemental figures and tables for "Multivariate distributions of behavioural, morphological, and ontogenetic traits in hybrids bring new insights into the divergence of sympatric Arctic charr morphs"

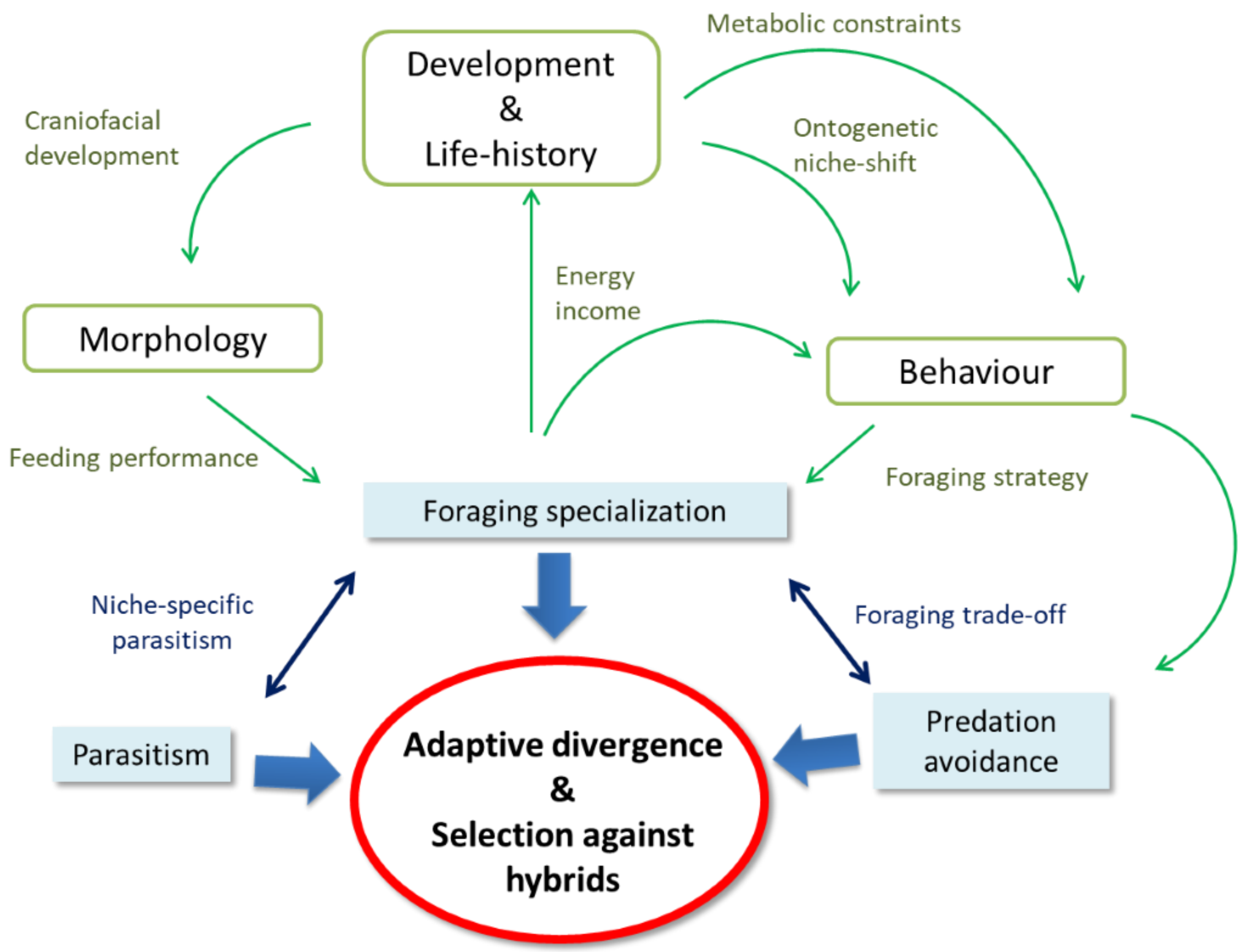


**Figure S1.** Dynamic framework illustrating in a non-exhaustive way how traits of different nature (green boxes) may be intertwined by functional constraints, which influences the response to different selective regimes (blue boxes) henceforth to process of speciation (red circle).

**Table S1.** Crossing design of the rearing experiment and number of individuals used for the personality tests. Type of cross: Female gamete x Male gamete. Parents were used for crosses only once (no split families). For the individual cells, the numbers of individuals on the left refer to the maximum number individuals at the start of the experiment. The numbers on the right refer to the number of individuals available with no missing data among all the sampling steps. One SBxPL family (19 individuals) from the "individual cells" setup hatched two weeks after the others and was no used for the analyses on trait covariance, growth, morphology and feeding behaviour.

| Set-up | Type of cross | Number of families | Number of individuals |
|---|---|---|---|
| Individual cells | PLxPL | 2 | 64 - 37 |
| | SBxSB | 2 | 42 -15 |
| | PLxSB | 3 | 75 - 23 |
| | SBxPL | 2 | 49 -18 |
| Group containers | PLxPL | 4 | 63 |
| | SBxSB | 3 | 32 |
| | PLxSB | 7 | 94 |
| | SBxPL | 3 | 25 |

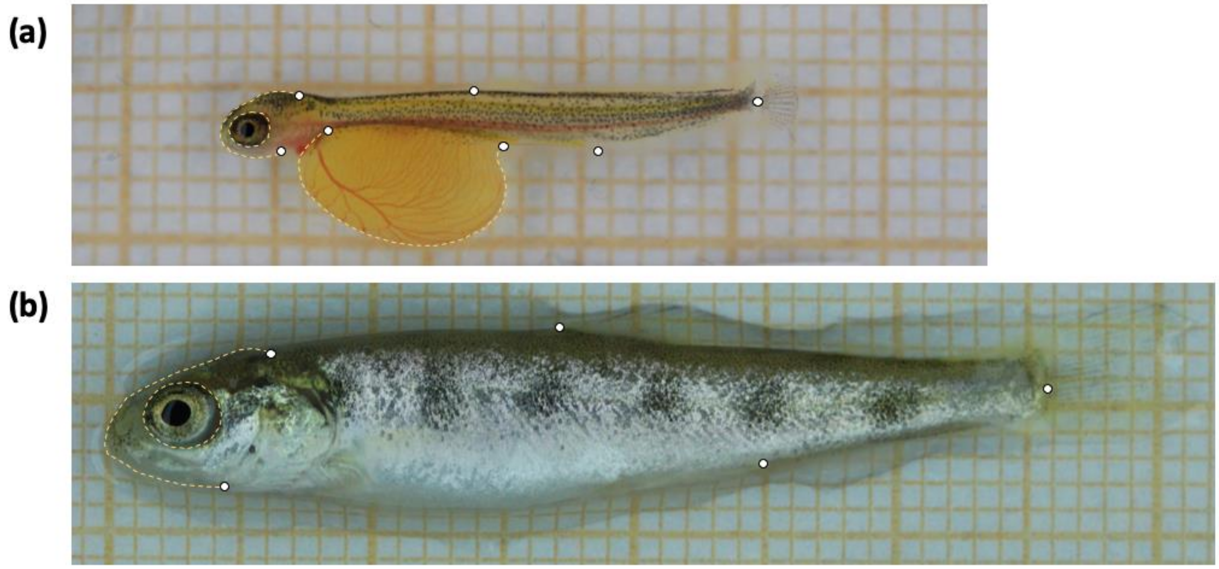


**Figure S2.** Landmarks used for the analyses of shape differences (a) on free-swimming embryos at hatching (D1: ca. 445 °C *d*ays) and (b) on actively feeding juveniles (D4: ca. 1100 °C *d*ays). The dashed curves depict the location of the Bezier curves used to extract the semi-landmarks. The specimens presented here are from the PLxPL offspring.

**Table S2.** List of the individuals from the individual cells that were discarded for the analyses.

| | Family of origin | Explanation |
|---|---|---|
| **Removed from the morphological analyses at hatching** | PLxPL | Identified as outlier in MANOVAs on body shape at hatching. Showed malformed mandibula and a unique shape of yolk sac on the photographs. |
| | SBxSB | Heavily malformed craniofacial morphology (“bulldog” face) |
| | SBxSB | Heavily malformed craniofacial morphology (“bulldog” face) |
| **Removed from the behavioural analyses** | PLxSB | Individual unable to correct its buoyancy |
| | SBxPL | Twisted spine constraining swimming activities |
| | PLxSB | Head infected by fongy |
| | SBxSB | Twisted spine constraining swimming activities |
| | SBxSB | Individual unable to correct its buoyancy |
| | PLxPL | Yolk sac not depleted and containing air |

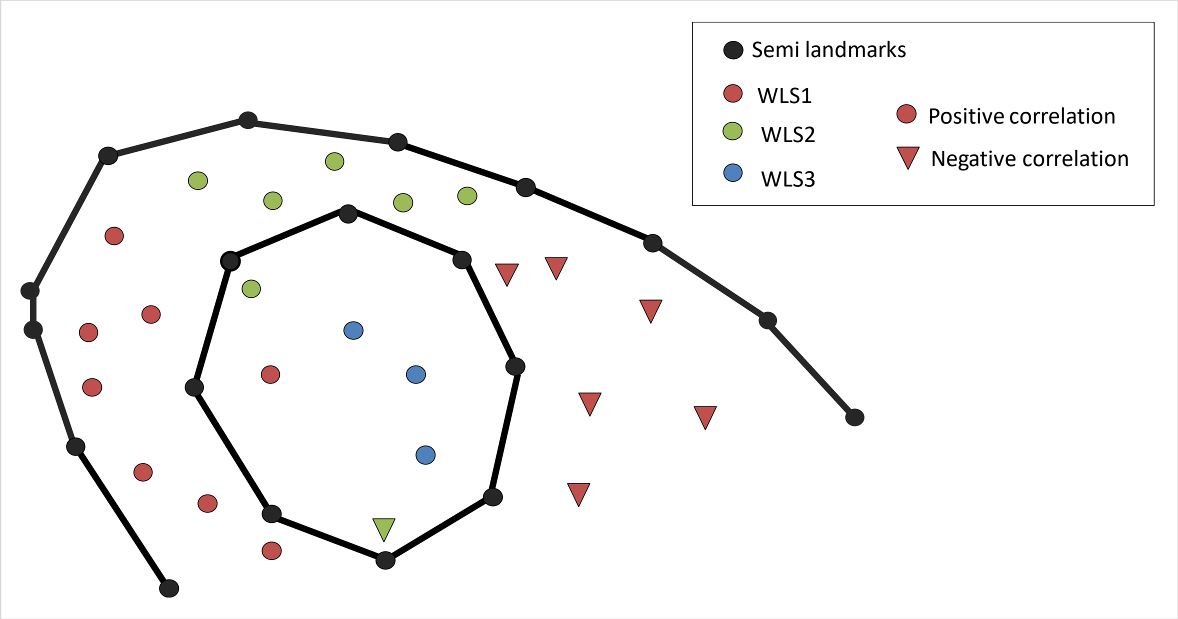


**Figure S3.** Locations of the Jacobian determinants loaded to the three latent variables obtained by Exploratory Factor Analysis (WLS1, WLS2 and WLS3). The shapes of the dots indicate whether the determinants are positively (points) or negatively (triangles) correlated to their respective latent variable.

**Table S3.** Specifications of the Generalized linear mixed-effect models run the analyses to the separate traits. For all model, burnin = 300 x number of iterations, thinning intervals = 10 x number of iterations.

| Trait | Fixed effects | Random variable | Model specificities | Number of iterations |
|---|---|---|---|---|
| **Growth** | | | | |
| Log standard length (average differences) | Cross type x Age | Family Individual | Random regression Second order polynomial regression | $6.5x10^4$ |
| Log standard length (within-group variation) | Age + Family | Individual | One model per type of cross | $1.3x10^6$ |
| **Yolk sac size at hatching and resorption** | | | | |
| Yolk sac area at hatching, Yolk sac area at hatching + 20 days (average differences) | Standard length + Cross type | Family Individual | Multi-response model | $3.9x10^6$ |
| Yolk sac area at hatching, Yolk sac area at hatching + 20 days (within-group variation) | Standard length Family | Individual | Multi-response model One model per type of cross | $4.6 x10^6$ |
| **Head shape** | | | | |
| WLS1 | Log standard length x Cross type | Family | | $1.3x10^4$ |
| WLS2 | Log standard length x Cross type | Family | | $3.9 x10^6$ |
| WLS3 | Log standard length x Cross type | Family | | $4.6 x10^6$ |
| **Feeding behaviour** | | | | |
| Age of exogeneous feeding | Cross type | Family | Binomial GLMM with logit link | $2.6 x10^5$ |
| Propensity to feed | Feeding trial x Cross type | Family Individual | | $3.9 x10^6$ |
| Log Latency to feed | Feeding trial x Cross type | Family Individual | | $2.6 x10^5$ |
| Total Number of feeding attempts | Feeding trial x Cross type | Family Individual | GLMM with log link | $5.7 x10^7$ |
| Number of feeding attempts (bottom) | Total Number of feeding attempts + Cross type | Family Individual | GLMM with log link | $1.0 x10^7$ |
| Number of feeding attempts (water column) | Total Number of feeding attempts + Cross type | Family Individual | GLMM with log link | $1.0 x10^7$ |
| Number of feeding attempts (surface) | Total Number of feeding attempts + Cross type | Family Individual | GLMM with log link | $1.0 x10^7$ |

**Table S4.** Posterior modes and 95%CrI (Credible Intervals) of the fixed effect of the quadratic regression on body length. Poly-1 & 2: First and second-order polynomial terms, respectively.

| | Posterior mode | 95% CrI |
|---|---|---|
| Intercept (PLxPL cross) | 3.09 | 2.97 - 3.19 |
| SBSB cross | -0.10 | -0.22 - 0.08 |
| Hybrid cross | -0.07 | -0.21 - 0.04 |
| Poly-1 | 6.14 | 5.89 - 6.38 |
| Poly-2 | -0.70 | -0.85 - -0.55 |
| Poly-1 SBxSB cross | -0.42 | -0.88 - -0.10 |
| Poly-1 Hybrid cross | -0.38 | -0.71 - -0.11 |
| Poly-2 SBxSB cross | -0.40 | -0.69 - -0.21 |
| Poly-2 Hybrid cross | -0.25 | -0.46 - -0.06 |

**Table S5.** Pairwise differences of the ontogenetic trajectories of head shape among types of crosses. Calculated differences, 95% Upper Confidence Limit , standardized scores and *p*-values are shown for three attributes of the trajectories (path length, angle and shape).

| | **Path length** | | | | **Angle** | | | | | **Shape** | | | |
|---|---|---|---|---|---|---|---|---|---|---|---|---|---|
| | *Δd* | UCLx | *Z* | *p* | *r* | Angle (°) | UCL | *Z* | *p* | *Δd* | UCL | *Z* | *p* |
| **PL-SB** | 0.01 | 0.03 | -0.28 | 0.53 | 0.99 | 7.03 | 9.59 | -0.72 | 0.76 | 0.14 | 0.22 | -0.34 | 0.62 |
| **PL-$F_1$** | 0.02 | 0.04 | 0.04 | 0.47 | 1.00 | 4.08 | 6.58 | -1.02 | 0.86 | 0.09 | 0.16 | -0.66 | 0.74 |
| **SB-$F_1$** | 0.03 | 0.05 | 0.40 | 0.35 | 1.00 | 3.79 | 6.81 | -1.02 | 0.86 | 0.15 | 0.22 | -0.33 | 0.63 |

**Table S6.** Procrustes variances and *p*-values of pairwise comparisons between types of crosses in the head shapes of the specimens reared individually, at D1 (hatching) and D3 (onset of first feeding).

| | **Procrustes variance** | | | ***p*** | | |
|---|---|---|---|---|---|---|
| | PL | SB | $F_1$ | PL-SB | PL-$F_1$ | SB-$F_1$ |
| **D1** | 0.00280 | 0.00372 | 0.00340 | 0.164 | 0.247 | 0.623 |
| **D3** | 0.00345 | 0.00249 | 0.00266 | 0.127 | 0.126 | 0.761 |

**Table S7.** Posterior estimates 95% Credible interval of the fixed effects of the three separates Generalized Linear Mixed Models on infinitesimal shape changes (WLS1, WLS2 and WLS3).

| | Effect | Posterior mode | 95% CrI | | |
|---|---|---|---|---|---|
| **WLS1** | | | | | |
| | Intercept (Cross PL) | -1.89 | -6.41 | - | 3.33 |
| | Log centroid size | 1.11 | -1.70 | - | 3.16 |
| | cross SB | 0.32 | -1.21 | - | 1.52 |
| | Cross PL | -0.41 | -1.31 | - | 0.88 |
| **WLS2** | | | | | |
| | Intercept (Cross PL) | 1.48 | -3.11 | - | 6.55 |
| | Log centroid size | -0.88 | -3.61 | - | 1.37 |
| | cross SB | 0.31 | -0.40 | - | 1.17 |
| | Cross PL | 0.39 | -0.23 | - | 1.04 |
| **WLS** | | | | | |
| | Intercept (Cross PL) | -1.76 | -5.60 | - | 2.84 |
| | Log centroid size | -0.23 | -1.59 | - | 2.59 |
| | cross SB | 0.57 | -0.19 | - | 1.27 |
| | Cross PL | 0.54 | -0.14 | - | 1.07 |

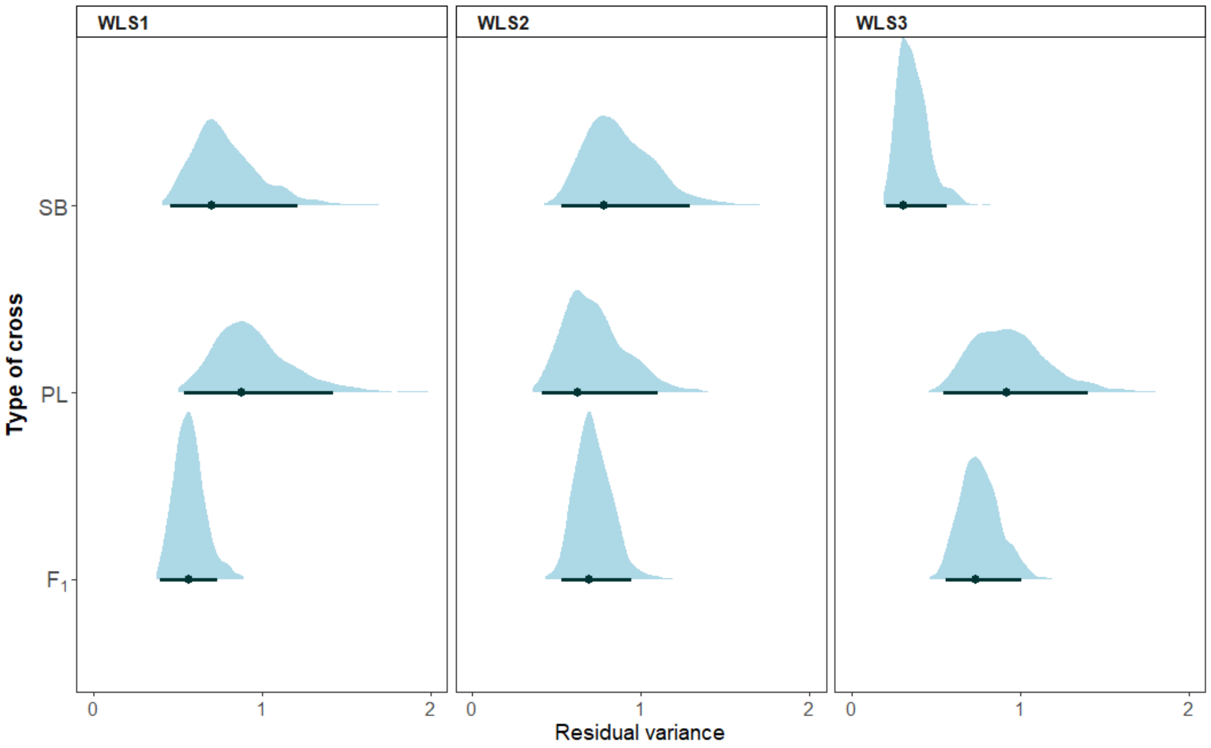


**Figure S4.** Posterior densities, Posterior modes and 95% Credible intervals of the three separate Generalized Linear Mixed Effects Models on latent variables of local shape changes (WLS1, WLS2 and WLS3).

**Table S8.** Posterior estimates of the Linear Mixed-effect Model on the age of exogeneous feeding (degree days). Categories of "cross" : cross SB = SBxSB offspring, cross PL = PLxPL offspring, cross $F_1$ = hybrids.

| | Effect | Posterior mode | 95% CrI |
|---|---|---|---|
| **Fixed effects** | Intercept (cross PL) | 651.7 | 642.5 - 662.1 |
| | cross SB | 5.1 | -10.4 - 17.5 |
| | cross $F_1$ | 3.3 | -9.1 - 15.0 |
| **Random effect variance** | Family | 0.5 | 0.0 - 146.9 |
| **Residuals** | cross PL | 228.3 | 154.6 - 377.6 |
| | cross SB | 323.1 | 226.1 - 626.9 |
| | cross $F_1$ | 253.5 | 200.7 - 387.4 |

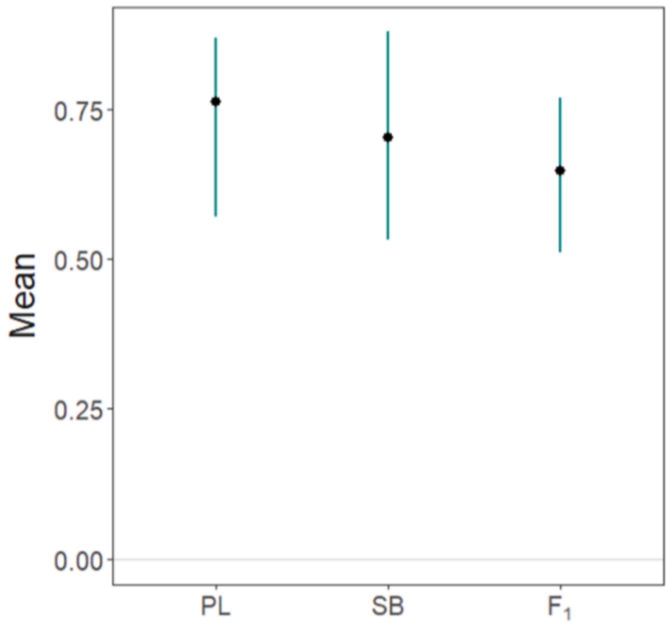


**Figure S5.** Posterior modes and 95% CrI of the mean propensity to start feeding during the observation trials. The estimates are obtained from a Generalised Linear Mixed-Effect Model relying on a Binomial distribution of link logit and are showed on the observed scale. Categories: SB = SBxSB offspring, PL = PLxPL offspring, $F_1$ = hybrids.

**Table S11.** Posterior estimates of the fixed effects of the Linear Mixed-Effect model on the latency of the focal individual to start feeding across the three observation trials on feeding behaviour (log seconds).

| | Posterior mode | 95% CrI | | |
|---|---|---|---|---|
| **Intercept (Cross PL : Trail 1)** | 3.12 | 642.48 | - | 662.14 |
| **Trial 2** | -0.38 | -10.41 | - | 17.52 |
| **Trial 3** | -0.44 | -9.11 | - | 15.01 |
| **Cross SB** | -0.51 | 642.48 | - | 662.14 |
| **Cross F1** | -0.55 | -10.41 | - | 17.52 |
| **Trial 2 : Cross SB** | 0.61 | -9.11 | - | 15.01 |
| **Trial 3 : Cross SB** | 1.10 | 642.48 | - | 662.14 |
| **Trial 2 : Cross F1** | 0.56 | -10.41 | - | 17.52 |
| **Trial 3 : Cross F1** | 0.37 | -9.11 | - | 15.01 |

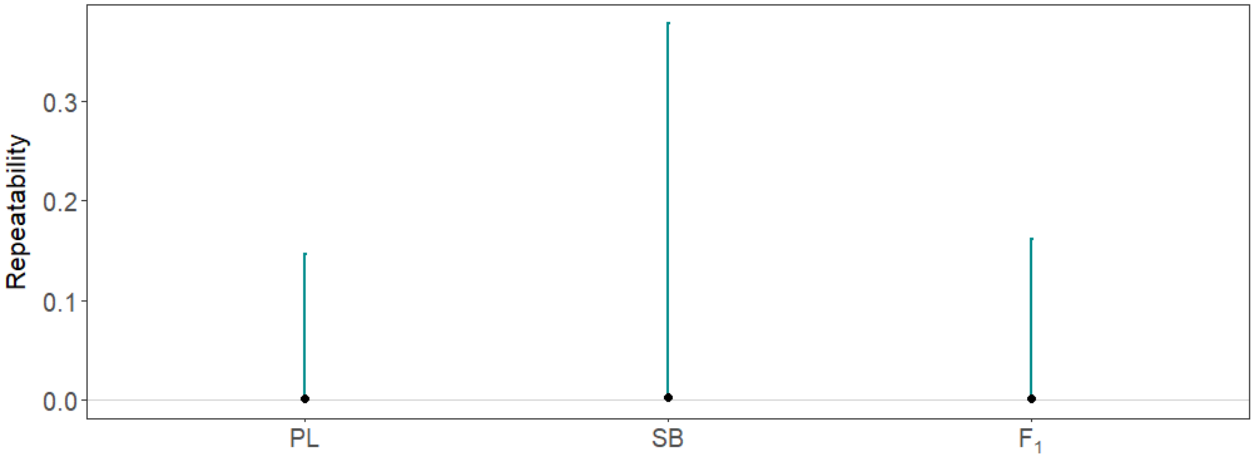


**Figure S4.** Posterior modes and 95% CrI of the repeatability of the latency to start feeding of the focal individuals observed during the feeding experiments, for each type of cross. Categories: SB = SBxSB offspring, PL = PLxPL offspring, $F_1$ = hybrids.

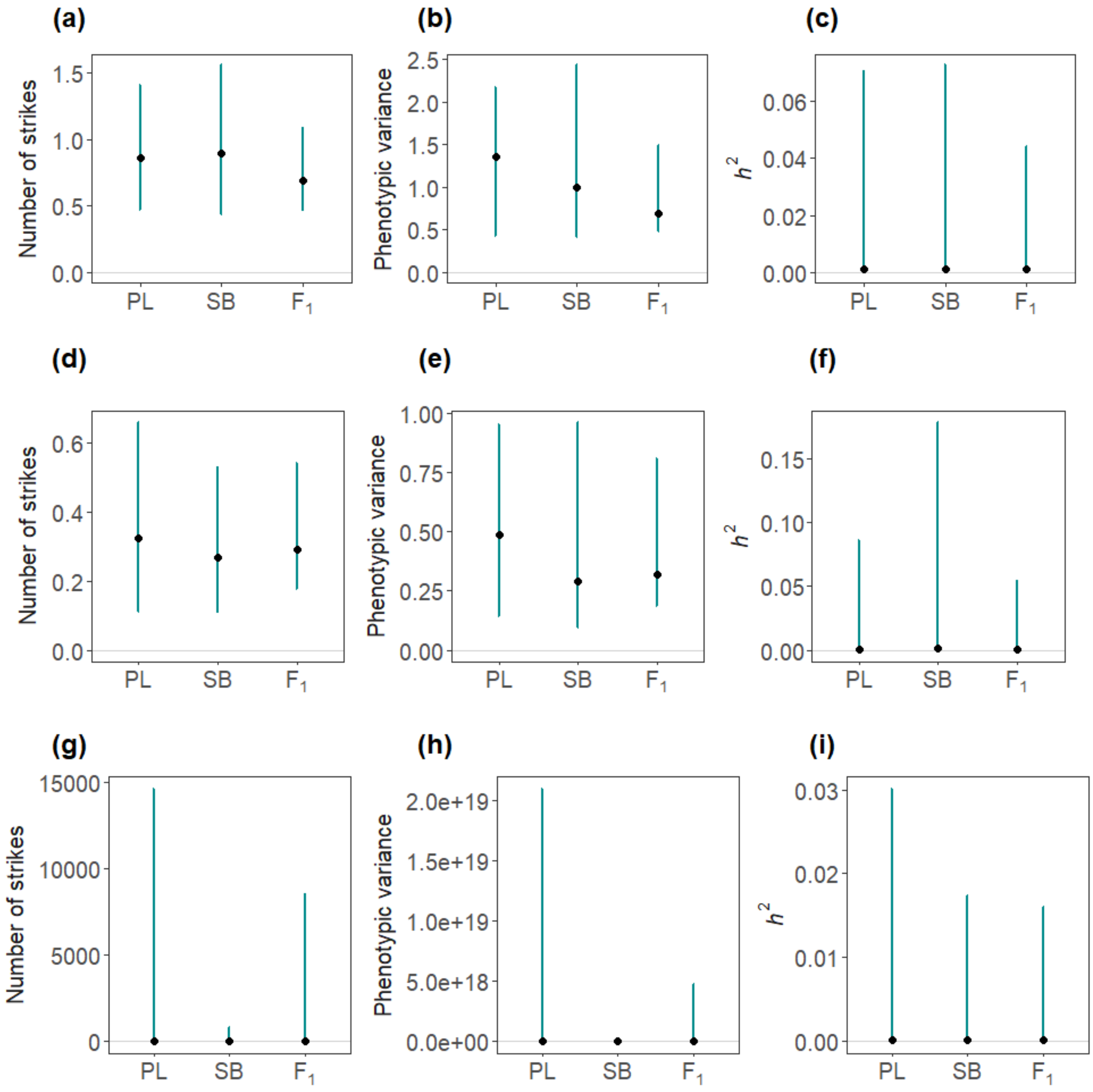


**Figure S5.** Posterior modes and 95% CrI of the of the numbers of feeding attempts recorded in each on the bottom of the cup (a-c) at mid-water (d-f) or on the surface (g-i) during the feeding trials. (a-g) Numbers of feeding attempts, (d-h) Total amount of within group variance, (c-i) heritability estimates($h^2$). Categories: SB = SBxSB offspring, PL = PLxPL offspring, $F_1$ = hybrids.

**Figure S6.** Posterior modes and 95% CrIs of the elements of the **P** matrices of each type of cross. Diagonal elements are variances and off-diagonal elements are correlations. Developmental time points D1 = hatching, D2 = 20 days post hatching, D3 = onset of exogeneous feeding, D4 = two months after the onset of exogeneous feeding. See text for details on the components of head shape.

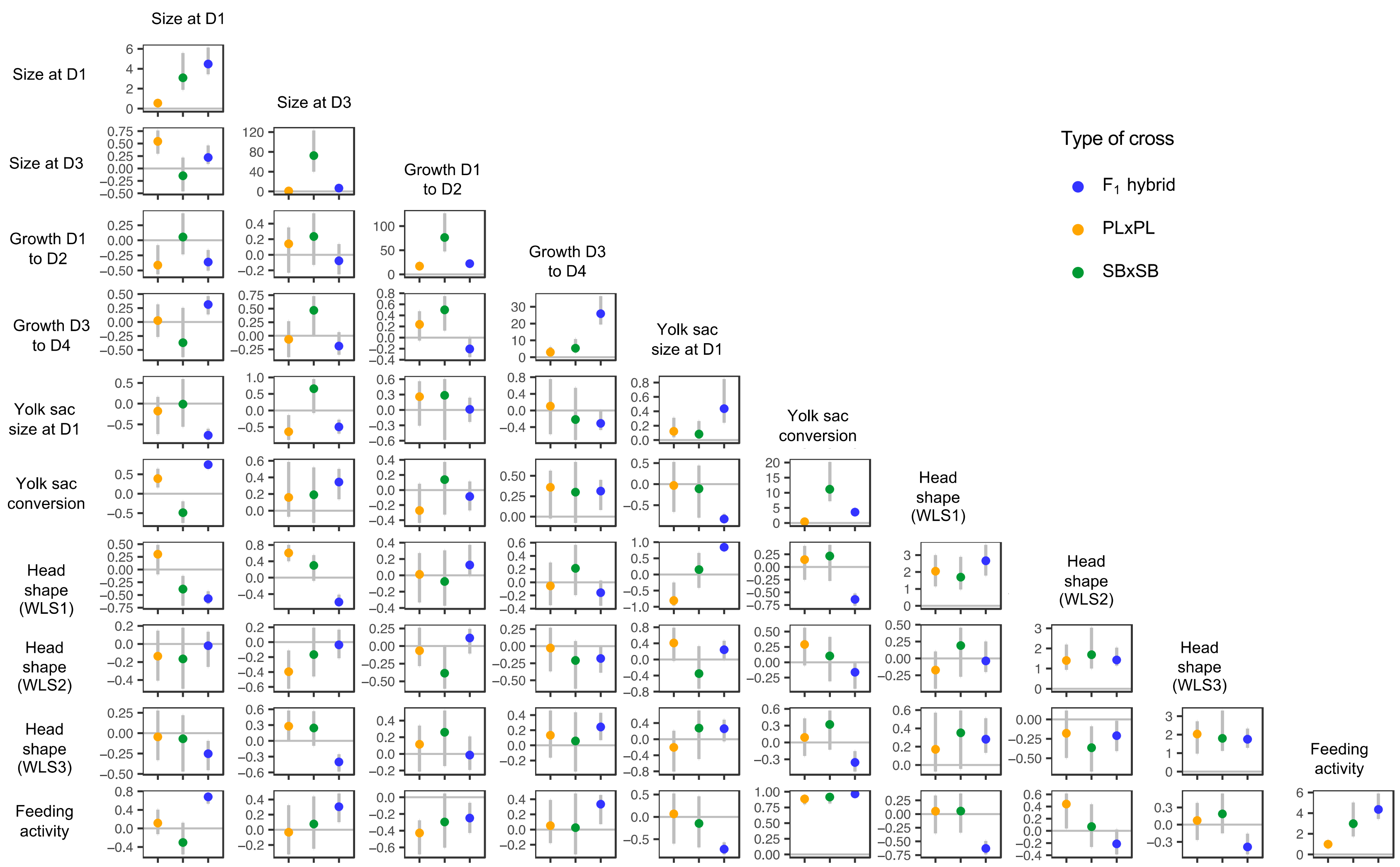

**Table S7.** Trait loadings of the first three eigenvectors of the **P** matrix of each type of cross (based on the posterior modes). Developmental time points D1 = hatching, D2 = 20 days post hatching, D3 = onset of exogeneous feeding, D4 = two months after the onset of exogeneous feeding. See text for details on the components of head shape.

| | **PLxPL** | | | **SBxSB** | | | **F1 Hybrids** | | |
|---|---|---|---|---|---|---|---|---|---|
| | **e1** | **e2** | **e3** | **e1** | **e2** | **e3** | **e1** | **e2** | **e3** |
| %Variance | 0.62 | 0.11 | 0.10 | 0.52 | 0.35 | 0.08 | 0.43 | 0.26 | 0.17 |
| Size at D1 | 0.05 | -0.15 | 0.08 | 0.00 | 0.06 | 0.23 | -0.19 | -0.08 | 0.36 |
| Size at D3 | -0.03 | -0.39 | 0.06 | 0.65 | -0.74 | 0.15 | 0.00 | -0.15 | 0.48 |
| Growth D1 to D2 | -0.98 | 0.01 | -0.10 | 0.74 | 0.66 | -0.02 | 0.54 | 0.80 | 0.26 |
| Growth D3 to D4 | -0.13 | 0.29 | 0.88 | 0.12 | 0.01 | -0.19 | -0.78 | 0.57 | -0.13 |
| Yolk sac age size at D1 | 0.03 | 0.02 | 0.18 | 0.09 | -0.10 | -0.84 | -0.14 | 0.02 | 0.44 |
| Yolk sac conversion | 0.12 | 0.09 | 0.20 | -0.03 | -0.08 | -0.41 | -0.18 | -0.04 | 0.43 |
| Head shape (WLS1) | -0.01 | -0.67 | 0.13 | 0.01 | -0.05 | -0.02 | 0.09 | 0.04 | -0.33 |
| Head shape (WLS2) | 0.03 | 0.29 | -0.03 | -0.05 | -0.02 | -0.06 | 0.04 | -0.02 | -0.01 |
| Head shape (WLS3) | -0.04 | -0.42 | 0.35 | 0.04 | -0.01 | -0.04 | -0.03 | 0.06 | -0.19 |
| Feeding_intensity | 0.00 | 0.16 | 0.00 | 0.01 | -0.02 | 0.03 | 0.04 | -0.01 | -0.18 |
